# An expanded GRAB sensor palette for multiplexed imaging of purinergic dynamics *in vivo*

**DOI:** 10.64898/2026.11.23.768484

**Authors:** Bohan Li, Yumo Li, Jing Wang, Zexiao Bai, Peiyao He, Zhixiong Ma, Yufei Hu, Fengru Li, Kaixin Cao, Weijian Teo, Chun Li, Kaikai He, Peilin Yang, Jing Wang, Yulong Li, Zhaofa Wu

## Abstract

Extracellular ATP and adenosine operate as tightly coupled purinergic signals, yet tracking both molecules simultaneously *in vivo* has been limited by the performance and spectral range of existing sensors. We describe an expanded set of genetically encoded GPCR-Activation-Based (GRAB) sensors for ATP and adenosine that together support multiplexed imaging of purinergic signaling in the intact brain. Systematic optimization of receptor scaffolds, membrane trafficking, and fluorophore coupling yielded next-generation green sensors (Ado2h, Ado2m, ATP2h, ATP2l) and spectrally separated red variants (rAdo1h, rATP1h), each combining high ligand specificity, genetically targetable expression, sub-second kinetics, and minimal engagement of endogenous GPCR pathways. This combination allows minimally invasive, dual-color imaging of purinergic activity *in vivo*. Using multiplexed fiber photometry and two-photon imaging, we tracked ATP and adenosine concurrently across several behavioral and pathological settings. ATP release preceded extracellular adenosine accumulation during sleep-wake transitions and neuronal activation, a pattern consistent with extracellular ATP hydrolysis as a substantial contributor to *in vivo* adenosine signaling. During epileptic seizures, paired imaging of extracellular ATP with extracellular adenosine, and of extracellular with intracellular adenosine, indicated distinct spatiotemporal dynamics across cellular compartments. This expanded sensor set offers a platform for multiplexed purinergic imaging and for probing ATP-adenosine coupling across physiological and pathological brain states.

## Introduction

Adenosine triphosphate (ATP) and adenosine (Ado) form a metabolically coupled purinergic signaling axis that regulates synaptic plasticity, sleep-wake regulation, motor control, and neuroprotection^1,2^. ATP and adenosine frequently pull neuronal excitability in opposite directions, yet their signaling remains coupled through extracellular metabolism: ATP released during neuronal activity is rapidly converted to adenosine by ectonucleotidases such as CD39 and CD73^3–5^. Purinergic signaling therefore depends less on the level of either molecule alone than on the balance between ATP release, extracellular enzymatic conversion of ATP to adenosine, and adenosine clearance. Disruption of this balance has been linked to epilepsy^6^, depression^7^, and sleep disorders^8^, which motivates resolving ATP and adenosine dynamics together, at matched spatial and temporal resolution, to understand how purinergic signaling shapes circuit function in health and disease^9,10^.

Recent studies have further revealed glial cells as active regulators of purinergic communication, demonstrating that ATP-adenosine gliotransmission contributes to the modulation of neuronal activity, behavioral states, and disease-associated circuit dysfunction^6,11–15^. Astroglial purinergic signaling has been implicated in norepinephrine-dependent regulation of behavioral state^13^, synaptic modulation^13^, and neuron-astrocyte interactions underlying depressive-like behaviors^11^. These findings emphasize the importance of monitoring purinergic dynamics across molecular species, cell types, and brain states, while underscoring the need for tools capable of multiplexed measurements *in vivo*.

Simultaneously resolving ATP and adenosine dynamics *in vivo* has proven difficult. Microdialysis and enzyme-based biosensors lack the temporal resolution, spatial precision, or multiplexing capacity needed to capture fast purinergic dynamics^9^. Genetically encoded GPCR-Activation-Based (GRAB) sensors have addressed similar limitations for other neuromodulators by combining high ligand specificity, genetically targetable expression, sub-second kinetics, and minimal engagement of native signaling cascades, together permitting chronic, minimally invasive, cell-type-specific recordings in behaving animals^16,17^. First-generation GRAB sensors for adenosine (Ado1.0)^18,19^ and ATP (ATP1.0)^20^ brought these advantages to purinergic signaling, but both operate in a single green channel with only moderate kinetics, which rules out multiplexed ATP-adenosine imaging and constrains analysis of their rapid coupling.

We set out to expand the GRAB sensor palette for multiplexed purinergic imaging *in vivo*. Systematic engineering of receptor scaffolds, membrane trafficking, and fluorophore coupling yielded next-generation green sensors and spectrally separated red variants for both ATP and adenosine. Their improved brightness, response speed, ligand specificity, and minimal engagement of endogenous GPCR pathways enabled simultaneous dual-color imaging across spatial and temporal scales. Using multiplexed fiber photometry and two-photon imaging in behaving mice, we asked whether extracellular ATP release precedes adenosine accumulation during sleep-wake transitions, locomotion, and neuronal stimulation, and whether ATP and adenosine show distinct extracellular and intracellular dynamics during epileptic seizures. This expanded GRAB sensor set provides a platform for multiplexed imaging of purinergic signaling in the intact brain.

## Results

### Engineering next-generation GRAB sensors for adenosine and ATP

To enable multiplexed imaging of purinergic signaling, we built a suite of improved green and red genetically encoded sensors for adenosine and ATP. Starting from the first-generation GRAB_Ado_ sensor Ado1.0, optimization of its circularly permuted EGFP (cpEGFP) module produced the intermediate Ado1.1 **(Extended Data Fig. 1a,b)**. Further tuning of the A_2A_ receptor (A_2A_R) scaffold, its N-terminal signal sequence, C-terminus, and residues involved in activation and signal transduction, yielded the high-affinity sensor Ado2h. Parallel engineering of the ligand-binding pocket produced the medium-affinity variant Ado2m **(Extended Data Fig. 1c**–**f)**. For a red-shifted counterpart, we inserted cpmApple into the third intracellular loop of A_2A_R and screened over 2,000 variants by optimizing the insertion site, linkers, and fluorescent protein module. This process yielded rAdo1h, which showed a ∼6-fold fluorescence increase upon adenosine application (**Extended Data Fig. 1g,h**). AlphaFold3-predicted structural models illustrate the architectures of these green and red GRAB-Ado sensors (**Fig. 1a**). All three sensors localized efficiently to the plasma membrane in HEK293T cells and showed ligand-evoked fluorescence increases (**Fig. 1b**). Their excitation and emission spectra were well separated, suited for dual-color imaging (**Fig. 1c**). Relative to Ado1.0, Ado2h, Ado2m, and rAdo1h all showed larger peak fluorescence responses (ΔF/F_0_) and improved signal-to-noise ratios (SNR) (**Fig. 1d,e**).

**Fig. 1:**
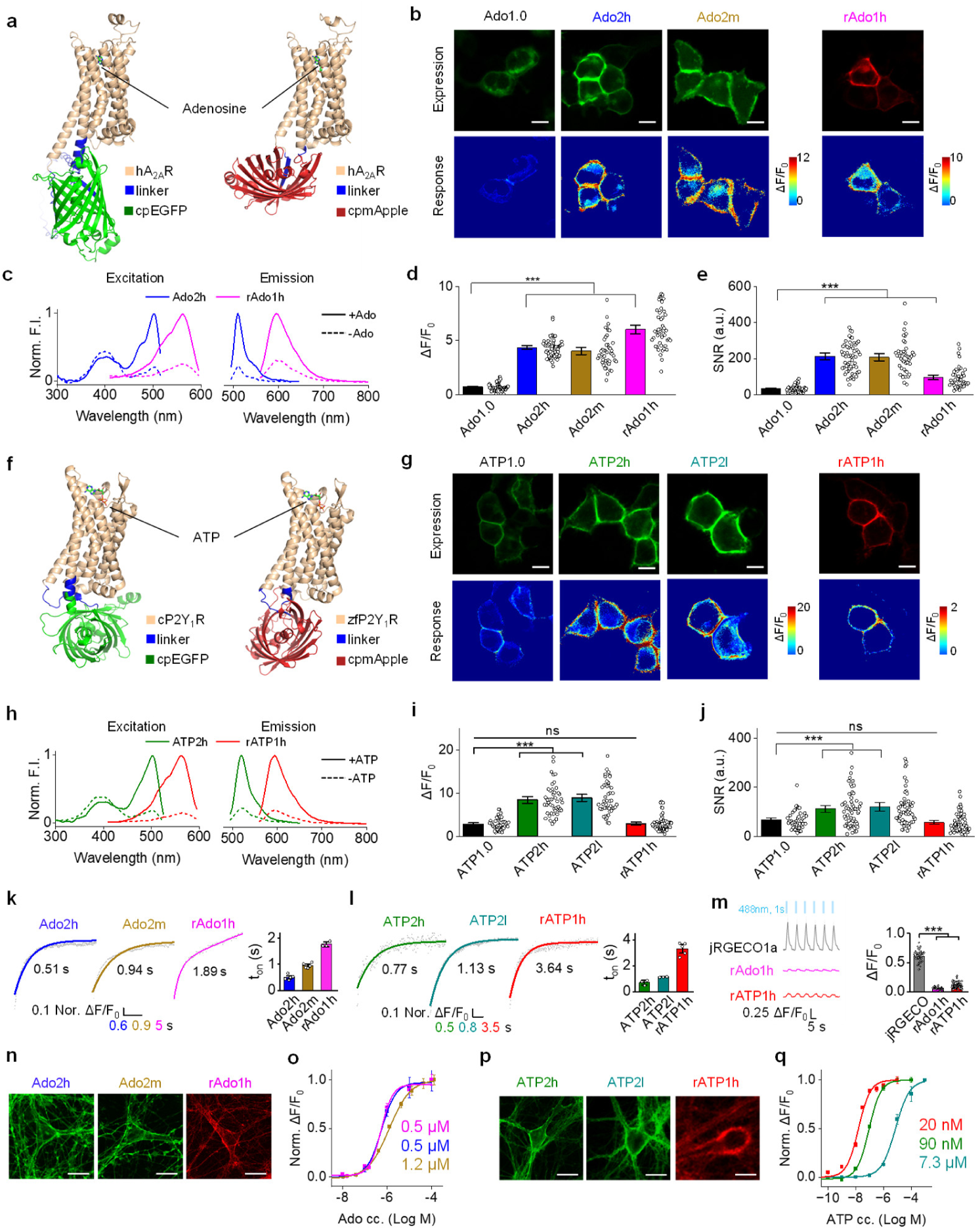
Development and performance of improved dual-color GRAB_Ado_ and GRAB_ATP_ sensors. **a**, AlphaFold3-predicted structures of the Ado2h and rAdo1h sensors, showing cpEGFP and cpmApple inserted into the third intracellular loop of A_2A_R, respectively. The N- and C-termini of the sensors are not shown. **b**, Representative fluorescence images of Ado1.0, Ado2h, Ado2m, and rAdo1h in HEK293T cells before and after 100 μM adenosine. Scale bar, 10 μm. **c**, One-photon excitation (left) and emission (right) spectra of Ado2h and rAdo1h with (solid lines) and without (dashed lines) adenosine. **d**, Peak responses of the indicated adenosine sensors. Bonferroni-adjusted *P* < 0.001 for Ado1.0 vs. Ado2h, Ado2m and rAdo1h; *n* = 45–60 cells from 3 independent experiments per sensor. **e**, Signal-to-noise ratios (SNRs; peak ΔF/F_0_ divided by the standard deviation of baseline ΔF/F_0_) for the indicated adenosine sensors. Bonferroni-adjusted *P* < 0.001 for Ado1.0 vs. Ado2h, Ado2m and rAdo1h; *n* = 45–60 cells from 3 independent experiments per sensor. **f**, AlphaFold3-predicted structures of ATP2h and rATP1h sensors, showing cpEGFP and cpmApple inserted into the third intracellular loop of P2Y_1_R, respectively. The N- and C-termini of the sensors are not shown. **g**, Representative fluorescence images of ATP1.0, ATP2h, ATP2l, and rATP1h in HEK293T cells before and after 100 μM ATP. Scale bar, 10 μm. **h**, One-photon excitation (left) and emission (right) spectra of ATP2h and rATP1h with and without ATP. **i**, Peak responses of the indicated ATP sensors. Bonferroni-adjusted *P* < 0.001 for ATP1.0 vs. ATP2h and ATP2l, and *P* = 0.886 for ATP1.0 vs. rATP1h; *n* = 45–60 cells from 3 independent experiments per sensor. **j**, SNRs for the indicated ATP sensors. Bonferroni-adjusted *P* = 0.0067 for ATP1.0 vs. ATP2h, *P* < 0.001 for ATP1.0 vs. ATP2l, and *P* > 0.999 for ATP1.0 vs. rATP1h; *n* = 45–60 cells from 3 independent experiments per sensor. **k**, Left, representative stopped-flow traces of Ado2h, Ado2m, and rAdo1h activation upon 100 μM adenosine. Right, quantified activation time constants; *n = 5, 5 and 3* traces, respectively. **l**, Left, representative stopped-flow traces of ATP2h and ATP2l activation upon 100 μM ATP and of rATP1h upon 30 μM ATP. Right, quantified activation time constants; *n = 3, 3 and 5* traces, respectively. **m**, Representative traces (left) and group summary of ΔF/F₀ (right) for the indicated sensors during 1 sec of 488 nm blue-light illumination (blue squares). Bonferroni-adjusted *P* < 0.001 for jRGECO1a vs. rAdo1h and rATP1h; *n* = 55, 50 and 100 cells from 3 independent experiments per sensor for jRGECO1a, rAdo1h and rATP1h, respectively. **n**, Representative fluorescence images of Ado2h, Ado2m, and rAdo1h in cultured neurons. Scale bar, 20 μm. **o**, Adenosine concentration-response curves for Ado2h, Ado2m, and rAdo1h; *n* = 45 neurons per sensor. **p**, Representative fluorescence images of ATP2h, ATP2l, and rATP1h in cultured neurons. Scale bar, 20 μm. **q**, ATP concentration-response curves for ATP2h, ATP2l, and rATP1h; *n* = 45 neurons per sensor from 3 independently prepared coverslips per sensor. Data are presented as mean ± s.e.m. Each data point represents one cell, neuron or stopped-flow trace, as indicated. For **d,e,i,j** and **m**, statistical significance was assessed by one-way ANOVA followed by Bonferroni’s multiple-comparisons test; ns, not significant; \*\**P* < 0.01, \*\*\**P* < 0.001.

We next optimized GRAB_ATP_ sensors. Because sequence variation can affect GPCR stability and conformational behavior^21–23^, we screened P2Y_1_ receptor (P2Y_1_R) orthologs for scaffolds potentially amenable to engineering. Replacing human P2Y_1_R with its chicken ortholog and optimizing the ligand pocket, cpEGFP module, and linkers gave rise to the high-affinity sensor ATP2h and the lower-affinity, ATP-preferring variant ATP2l (**Extended Data Fig. 1i,j**). In parallel, introducing cpmApple into zebra finch P2Y_1_R yielded the red sensor rATP1h (**Extended Data Fig. 1m,n**); AlphaFold3-predicted structures depict the sensor architectures (**Fig. 1f**). All three ATP sensors localized to the plasma membrane and showed ATP-evoked fluorescence increases in HEK293T cells (**Fig. 1g**). The green sensors ATP2h and ATP2l and the red sensor rATP1h had well-separated excitation and emission spectra (**Fig. 1h**). Dose-response measurements in cultured neurons further distinguished the two green variants. ATP2h detected ATP and ADP with similarly high affinity (EC_50_, 110 and 105 nM, respectively), showing minimal preference for ATP. By contrast, ATP2l had lower affinity but preferentially responded to ATP over ADP (EC_50_, 5.1 and 14 μM, respectively), with a reduced maximal response to ADP (**Extended Data Fig. 1k**). ATP2h and ATP2l showed larger peak ΔF/F_0_ and higher SNRs than the previously reported ATP1.0^20^ (**Extended Data Fig. 1l**), and rATP1h showed a red fluorescence response with an SNR comparable to that of ATP1.0 (**Fig. 1i,j**). Full-length sensor sequences are provided in **Extended Data Fige. 2–4**.

Because transient purinergic signals require rapid detection, we next measured activation kinetics. Ado2h, Ado2m, and rAdo1h activated on a subsecond-to-second timescale (**Fig. 1k**), as did ATP2h, ATP2l, and rATP1h (**Fig. 1l**). We also tested whether the red-shifted sensors were prone to blue-light-induced photoactivation, a known artifact of the cpmApple-based red calcium indicator jRGECO1a^24,25^. Brief 488-nm pulses produced smaller fluorescence changes in rAdo1h and rATP1h than in jRGECO1a, suggesting reduced sensitivity to blue-light illumination (**Fig. 1m**).

In cultured neurons, all six sensors trafficked efficiently to the membrane (**Fig. 1n,p**). Ado2h, Ado2m, and rAdo1h had half-maximal effective concentrations (EC_50_) of 0.5 μM, 1.2 μM, and 0.5 μM, respectively (**Fig. 1o**), while ATP2h, ATP2l, and rATP1h had EC_50_ values of 90 nM, 7.3 μM, and 20 nM (**Fig. 1q**). Ligand-insensitive control sensors showed no detectable response to their cognate ligands (**Extended Data Fig. 5a,b**).

Finally, we benchmarked the new sensors against existing purine sensors in HEK293T cells. With 100 μM adenosine, Ado2h showed a 4.1-fold increase in relative brightness, versus 2.5-fold for AdoLight^26^ and 1.8-fold for Ado1.0 (**Extended Data Fig. 5c**). With 1 mM ATP, ATP2l showed a 4.56-fold brightness increase, whereas iATPSnFR2.0-A119L^27^, membrane-displayed via a PDGFR transmembrane (TM) domain, showed only a 1.05-fold increase under identical conditions (**Extended Data Fig. 5d,e**). Together, these comparisons support the larger ligand-evoked fluorescence responses of the new sensor set.

### Specificity, pharmacology, and dual-color performance of the sensor suite

Multiplexed purinergic imaging requires sensors with ligand discrimination, minimal perturbation of native signaling, and clean spectral separation, so we next characterized these properties directly. Because extracellular ATP is sequentially hydrolyzed to adenosine by ectonucleotidases, the two signals are metabolically linked, making ligand discrimination a design requirement (**Fig. 2a**). In HEK293T cells, Ado2h, Ado2m, and rAdo1h responded to adenosine with negligible responses to other purines or neurotransmitters, and adenosine-evoked signals were largely blocked by the A_2A_R antagonist SCH-58261, consistent with receptor-specific activation (**Fig. 2b1**). ATP2h, ATP2l, and rATP1h responded strongly to ATP, and these responses were abolished by the P2Y_1_R antagonist MRS-2500 (**Fig. 2b2**). ADP is a known P2Y_1_R agonist^28^, and among the three ATP sensors, ATP2l showed the greatest preference for ATP over ADP, limiting potential cross-reactivity in settings where both nucleotides are present.

**Fig. 2:**
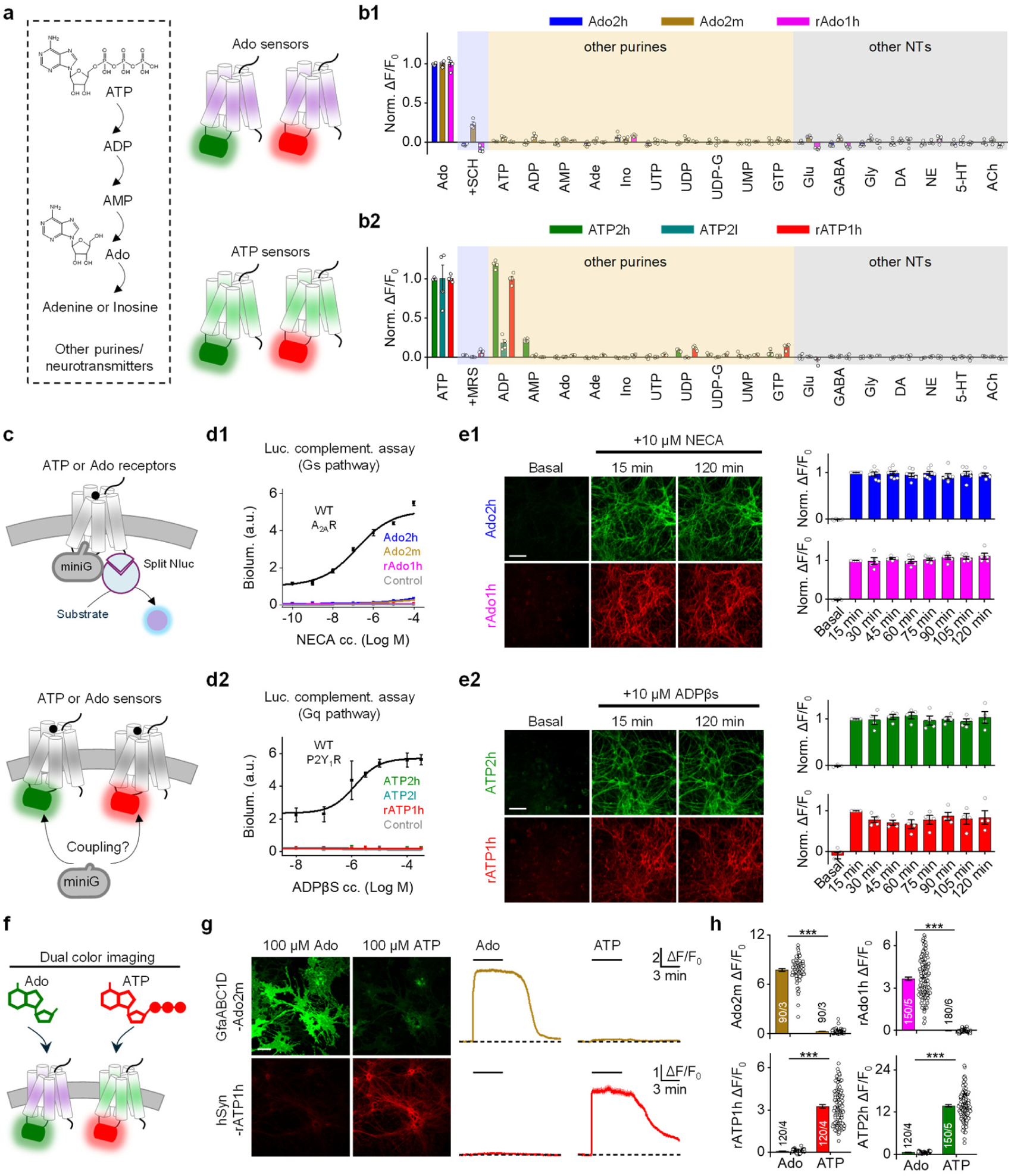
Characterization of GRAB_Ado_ and GRAB_ATP_ sensors in cultured cells. **a,** Schematic of purinergic signaling pathways. **b,** Pharmacological specificity of GRAB_Ado_ (top) and GRAB_ATP_ (bottom) sensors in HEK293T cells. **b1**, Ado sensor responses to purinergic ligands and neurochemicals (100 μM), with antagonists at 50 μM. **b2**, ATP sensor responses to the same ligand panel at 10 μM, with antagonists at 30 μM. *n* = 3–4 biological replicates (wells), ∼400 cells analyzed per well. SCH-58261 (SCH), A_2A_R antagonist; MRS-2500 (MRS), P2Y_1_R antagonist; ATP, adenosine triphosphate; ADP, adenosine diphosphate; AMP, adenosine monophosphate; Ado, adenosine; Ade, adenine; Ino, inosine; UTP, uridine triphosphate; UDP, uridine diphosphate; UDP-G, uridine diphosphate-glucose; UMP, uridine monophosphate; GTP, guanosine triphosphate; Glu, glutamate; GABA, γ-aminobutyric acid; Gly, glycine; DA, dopamine; NE, norepinephrine; 5-HT, serotonin; ACh, acetylcholine. **c**, Schematic of the miniG-based split-luciferase complementation assay used to assess receptor-G-protein coupling. **d1**, NECA concentration-response curves (miniGs-based complementation) for Ado2h, Ado2m, rAdo1h, wild-type A_2A_R, and control. **d2**, ADPβS concentration-response curves (miniGq-based complementation) for ATP2h, ATP2l, rATP1h, wild-type P2Y_1_R, and control. Cells expressing miniGs or miniGq alone served as controls; *n* = 3 independent cultures. **e**, Normalized sensor fluorescence over 2 hours of agonist exposure in cultured neurons. **e1**, Representative images (left) and fluorescence time course (right) for Ado2h and rAdo1h during 10 μM NECA application. **e2**, Representative images (left) and fluorescence time course (right) for ATP2h and rATP1h during 10 μM ADPβS application. *n* = 5–7 wells per condition, ∼100 neurons analyzed per well. Scale bar, 100 µm. **f**, Schematic of the dual-color imaging setup. **g**, Dual-color imaging in a co-culture expressing Ado2m in astrocytes and rATP1h in neurons. Left, representative fluorescence images; scale bar, 50 µm. Right, simultaneous Ado2m and rATP1h fluorescence traces during sequential application of 100 µM Ado and 100 µM ATP. **h**, Cross-reactivity of Ado and ATP sensors in astrocytes and neurons, shown as peak ΔF/F_0_ responses to 100 µM Ado or 100 µM ATP. *P <* 0.001 (Ado vs. ATP). Numbers on bars indicate ROIs/independent cultures. Data are presented as mean ± s.e.m. Each data point represents one sample. Student’s *t*-test; \*\*\**P <* 0.001.

We then asked whether the engineered receptors retained residual G-protein coupling, since unintended signaling could alter cell physiology. MiniG-based split-luciferase complementation assays^29^ showed negligible Gs activation for Ado2h, Ado2m, and rAdo1h relative to wild-type A_2A_R, and negligible Gq activation for ATP2h, ATP2l, and rATP1h relative to wild-type P2Y_1_R (**Fig. 2c,d1,d2**). Prolonged exposure to saturating concentrations of the stable agonists NECA (A_2A_R) or ADPβS (P2Y_1_R) caused no detectable loss of membrane fluorescence over 120 minutes (**Fig. 2e1,e2**), indicating little agonist-induced internalization. These properties support the use of the sensors for extended recordings without confounding signaling or trafficking effects.

Finally, we tested spectral and ligand specificity under dual-color conditions in neuron-astrocyte co-cultures (**Fig. 2f**). Expressing the green adenosine sensor Ado2m in astrocytes alongside the red ATP sensor rATP1h in neurons showed that Ado2m responded selectively to adenosine and rATP1h selectively to ATP, with no detectable cross-reactivity (**Fig. 2g**). The reciprocal configuration, green ATP2h in neurons, red rAdo1h in astrocytes, showed the same clean separation, with quantitative analysis showing minimal channel crosstalk (**Fig. 2h**).

Together, these results define a spectrally separable sensor palette with high ligand specificity, negligible G-protein coupling, and limited agonist-induced internalization, providing a foundation for simultaneous, cell-type-resolved imaging of adenosine and ATP dynamics in living neural circuits.

### Ado2h reports adenosine dynamics across sleep-wake cycles and acute stress

To evaluate Ado2h *in vivo*, we focused on the basal forebrain (BF), where adenosine released from VGLUT2+ neurons contributes to sleep regulation^18^. We first asked whether Ado2h could detect optogenetically evoked adenosine release. Adeno-associated viruses (AAVs) expressing Cre-dependent ChrimsonR^30^ were injected into the BF of VGLUT2-IRES-Cre mice together with Ado2h (**Extended Data Fig. 6a**). Light stimulation at 20 Hz produced time-locked increases in Ado2h fluorescence, with longer stimulation trains yielding progressively larger responses (**Extended Data Fig. 6b,c**). Pharmacological and mutational controls supported the specificity of the Ado2h signal. The adenosine kinase inhibitor 5-iodotubercidin (5-ITU) gradually increased basal fluorescence, enhanced stimulation-evoked responses and slowed their decay (**Extended Data Fig. 6b–d**). JMF1907, which acts as both an A_2A_R agonist and an equilibrative nucleoside transporter 1 blocker, also altered Ado2h signal amplitude and kinetics (**Extended Data Fig. 6e–g**). Conversely, the A_2A_R antagonist SCH-58261 suppressed responses (**Extended Data Fig. 6e–g**), and the sensor mutant Adomut showed no detectable signal (**Extended Data Fig. 6h–i**). Together, these results support the use of Ado2h to report endogenous adenosine dynamics in the BF.

We then compared Ado2h directly against the first-generation sensor Ado1.0^18^. To limit inter-animal variability, we injected AAVs expressing each sensor under the GfaABC1D promoter into opposite hemispheres of the BF within the same animal, allowing simultaneous recording (**Fig. 3a**). As previously reported for Ado1.0^18^, both sensors registered higher adenosine levels during wakefulness and REM sleep than during NREM sleep. Ado2h showed a significantly larger response amplitude during REM sleep and a faster off-rate during state transitions than Ado1.0 (**Fig. 3b**–**f**). Cross-correlation analysis showed weaker temporal agreement between the two sensors specifically during NREM sleep (**Extended Data Fig. 6j**), likely reflecting Ado2h’s ability to resolve spontaneous adenosine release events that Ado1.0 misses (**Fig. 3g,h**). The A_2A_R antagonist SCH-58261 reduced all Ado2h responses, supporting the specificity of the signal (**Fig. 3i**–**k**).

**Fig. 3:**
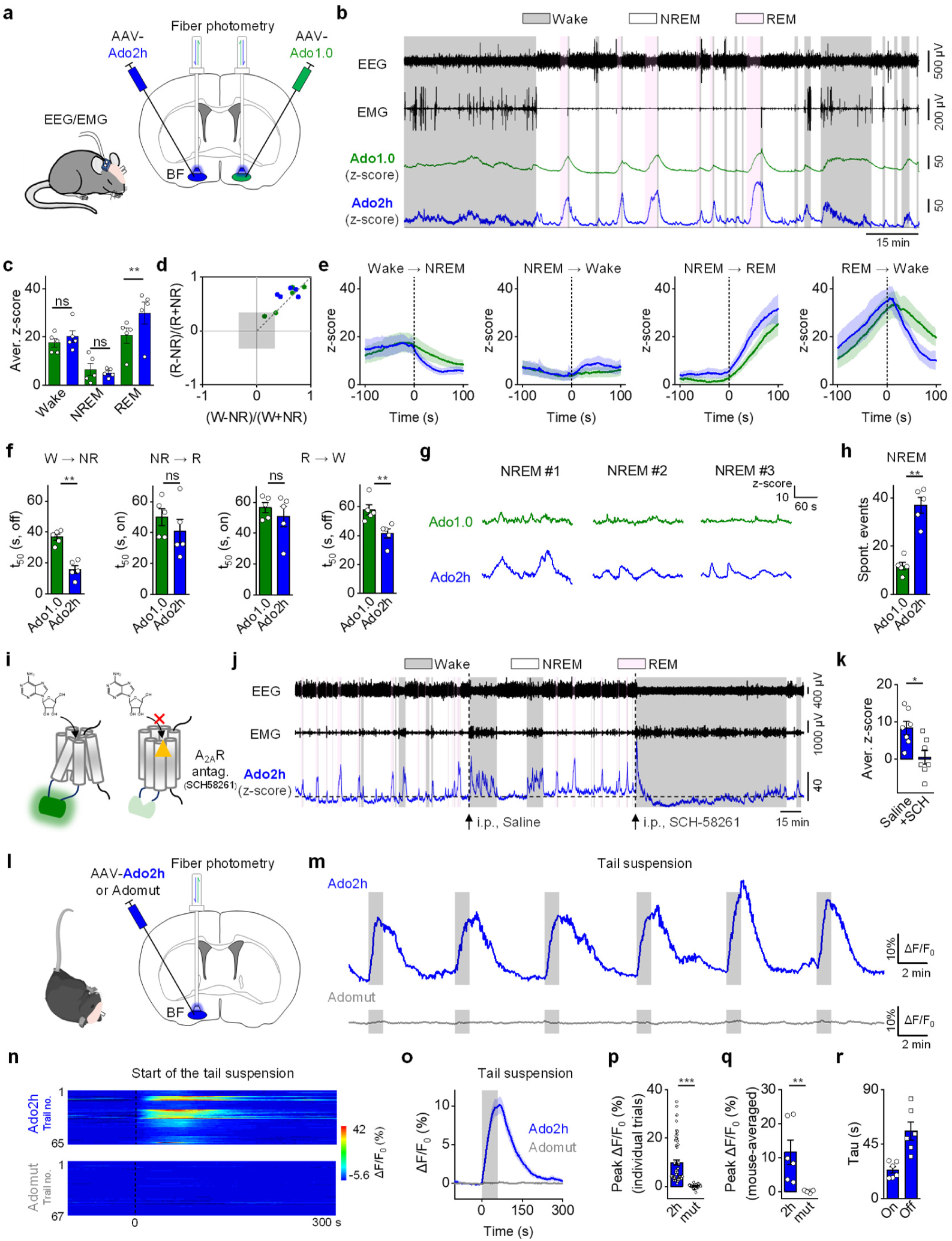
*In vivo* characterization of GRAB_Ado_ sensors in the mouse basal forebrain during sleep-wake cycles and acute stress. **a,** Schematic of fiber photometry recording of Ado1.0 and Ado2h in the basal forebrain (BF). **b**, Representative simultaneous EEG, EMG, and z-score traces for Ado1.0 (green) and Ado2h (blue) across sleep-wake cycles. Shading: wakefulness (grey), REM sleep (pink), NREM sleep (unshaded). W, wake; NR, NREM sleep; R, REM sleep. **c**, Summary of Ado1.0 and Ado2h fluorescence (z-score) by vigilance state. Two-way repeated-measures ANOVA revealed a significant sensor-by-vigilance-state interaction (*P =* 0.010). Post hoc comparisons showed no significant difference between Ado1.0 and Ado2h during wakefulness (*P =* 0.072) or NREM sleep (*P >* 0.99), but a significant difference during REM sleep (*P =* 0.0013). Each symbol represents one recording session. *n =* 5 sessions from 3 mice in c–f. **d**, Normalized signal modulation, (REM−NREM)/(REM+NREM) and (Wake−NREM)/(Wake+NREM), for each sensor. Grey box marks values under a twofold difference. **e**, Time-aligned average fluorescence for Ado1.0 and Ado2h centered on sleep-wake transitions (time 0, dashed line). **f**, Statistical comparison of the signal changes in e. *P =* 0.008 (W → NR), *P* = 0.31 (NR → R), *P =* 0.55 for ton and *P* = 0.008 for toff (R → W). **g**, Representative Ado1.0 and Ado2h traces across three NREM episodes (NREM #1–3). **h**, Number of spontaneous fluorescence events during NREM sleep for each sensor. *P* = 0.008; *n* = 5 sessions from 3 mice. **i**, Schematic of Ado signaling and the mechanism of the A_2A_R antagonist SCH-58261. **j**, Representative Ado2h fluorescence (ΔF/F_0_) in the BF across sleep-wake cycles under baseline, saline (i.p.), and SCH-58261 (i.p., 10 mg/kg) conditions. Arrow marks injection time. **k**, Summary of Ado2h fluorescence changes after saline or SCH-58261. *P =* 0.02; *n* = 7 mice per group. **l**, Schematic of recording Ado2h and the non-responsive control Adomut in the BF during tail-suspension stress. **m**, Representative ΔF/F_0_ traces for Ado2h (blue) and Adomut (grey) before, during (grey bar), and after tail suspension. **n**, Heatmaps of all trials for Ado2h (65 trials, 6 mice) and Adomut (67 trials, 3 mice) aligned to tail-suspension onset. **o**, Average ΔF/F_0_ responses for Ado2h and Adomut across all trials, aligned to tail suspension (grey bar). **p**, Summary of peak ΔF/F_0_ across all trials (65 trials, 6 mice for Ado2h; 67 trials, 3 mice for Adomut). *P <* 0.001. **q**, Summary of peak ΔF/F_0_ averaged per mouse. *P =* 0.002; *n* = 6 mice for Ado2h, *n* = 3 mice for Adomut. **r**, Kinetics of the Ado2h response to tail suspension: rise (Tau-on) and decay (Tau-off) time constants; *n* = 6 mice. Data are presented as mean ± s.e.m. In **c**, data were analyzed using two-way repeated-measures ANOVA with sensor and vigilance state as within-session factors, followed by Bonferroni-corrected comparisons between sensors within each vigilance state. Other indicated comparisons were performed using two-sided Mann-Whitney *U* tests; ns, not significant, \**P <* 0.05, \*\**P <* 0.01, \*\*\**P <* 0.001.

Adenosine signaling has also been implicated in stress and antidepressant responses: deep brain stimulation^31^, ketamine, and electroconvulsive therapy^7^ can each trigger adenosine release, and enhanced A_1_R signaling produces antidepressant-like effects^32^. Taking advantage of Ado2h’s faster kinetics and larger dynamic range, we expressed it in the BF and subjected mice to tail-suspension stress, observing a reproducible, time-locked rise in Ado2h fluorescence during each stress episode (**Fig. 3l**–**r**). The Adomut control showed no response under identical conditions (**Fig. 3m**–**r**), indicating that the signal reflects genuine adenosine release and that acute stress rapidly engages endogenous adenosine signaling in the BF.

Together, these experiments show that Ado2h resolves endogenous adenosine dynamics across sleep-wake states and acute stress with high sensitivity and specificity, capturing rapid adenosine transients tied to behavioral state transitions and stress responses. The improved kinetics and SNR measured *in vitro* are consistent with improved *in vivo* detection of spontaneous and evoked adenosine signals.

### Optogenetically evoked adenosine and ATP release *in vivo*

To evaluate the red-shifted adenosine sensor rAdo1h *in vivo*, we co-expressed rAdo1h, Cre-dependent GCaMP6s, and Cre-dependent ChrimsonR in hippocampal CA1 of CaMKIIα-Cre mice (**Fig. 4a,b**). Optogenetic activation of CA1 pyramidal neurons produced time-locked increases in both GCaMP6s and rAdo1h fluorescence (**Fig. 4c,e**). Stimulation at 20 Hz for 10 s induced an epileptiform afterdischarge, as indicated by elevated electroencephalogram (EEG) power (**Fig. 4d**). During the afterdischarge, both calcium and adenosine signals showed a biphasic profile with a pronounced second phase (**Fig. 4e**), consistent with the calcium dynamics reported previously^33^. rAdo1h displayed a higher peak ΔF/F_0_ than GCaMP6s, indicating a larger dynamic response during epileptiform activity.

**Fig. 4:**
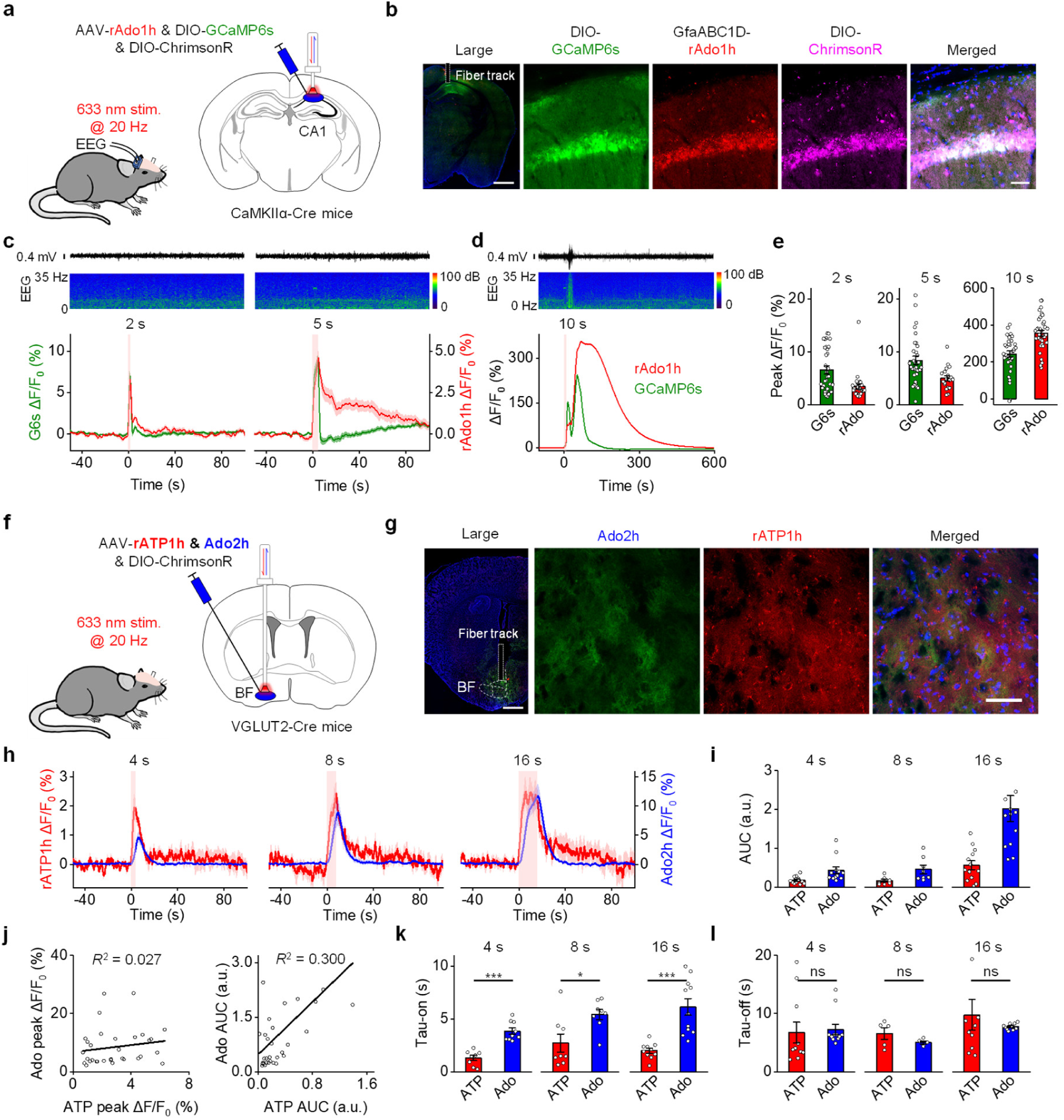
Detection of optogenetically evoked ATP and adenosine release in freely moving mice. **a,** Experimental design for co-expressing rAdo1h, GCaMP6s, and ChrimsonR with fiber photometry recording in hippocampal CA1 of CaMKIIα-Cre mice. **b**, Representative histology showing rAdo1h (red), GCaMP6s (green), and ChrimsonR (purple) expression in CA1; nuclei stained with DAPI (blue). Dashed white rectangle marks the optical-fiber track. Scale bars, 500 µm (left), 20 µm (right). **c**, Representative EEG traces and spectrograms (top) and trial-averaged rAdo1h and GCaMP6s fluorescence (bottom) during 2- and 5-s optical stimulation at 20 Hz. **d**, Representative EEG trace and spectrogram (top) and trial-averaged rAdo1h and GCaMP6s responses (bottom) during 10-s optical stimulation, with seizure stages indicated. **e**, Peak ΔF/F_0_ responses of rAdo1h and GCaMP6s for the indicated stimulation durations. Each point is one trial; *n* = 19–35 trials from 3 mice. **f**, Experimental design for co-expressing rATP1h, Ado2h, and ChrimsonR with fiber photometry recording in the BF of VGLUT2-Cre mice. **g**, Representative histology showing rATP1h (red) and Ado2h (green) expression in the BF; nuclei stained with DAPI (blue). Dashed white rectangle marks the optical-fiber track. Scale bars, 500 µm (left), 20 µm (right). **h**, Trial-averaged rATP1h (red) and Ado2h (blue) fluorescence during 4-, 8-, and 16-s optical stimulation; red shading marks the stimulation period. **i**, Area under the curve (AUC) for the responses in h; *n* = 8–13 trials from 3 mice per stimulation duration. **j**, Trial-level relationships between rATP1h and Ado2h peak ΔF/F_0_ (left) and AUC (right), pooled across 4-, 8-, and 16-s stimulation. For peak ΔF/F_0_: Pearson’s *r*(30) = 0.163, two-sided *P* = 0.373, 95% CI [−0.197, 0.484]. For AUC: Pearson’s *r*(30) = 0.547, two-sided *P* = 0.00118, 95% CI [0.246, 0.753]. *n* = 32 trials from 3 mice; each point is one trial, black lines show linear fits with *R*^2^ values indicated. **k**, Rise time constants (Tau-on) of rATP1h and Ado2h responses. *P* < 0.001 (4 s), *P* = 0.02 (8 s), and *P* < 0.001 (16 s). **l**, Decay time constants (Tau-off) of rATP1h and Ado2h responses. *P* = 0.07 (4 s), *P* = 0.42 (8 s), *P* = 0.85 (16 s). Data are presented as mean ± s.e.m. Each data point represents one sample. Mann-Whitney *U* test; ns, not significant; \**P* < 0.05, \*\*\**P* < 0.001.

Extracellular ATP is rapidly hydrolyzed to adenosine by ectonucleotidases, yet the two purines signal through distinct receptor families that coordinate separate physiological processes^34^. To compare their spatiotemporal dynamics directly, we co-expressed the red ATP sensor rATP1h and the green adenosine sensor Ado2h in the BF and optogenetically stimulated VGLUT2+ neurons (**Fig. 4f,g**). Lengthening the 20-Hz stimulation train progressively increased the area under the curve for both signals (**Fig. 4h–j**), with rATP1h showing faster onset kinetics than Ado2h while their decay kinetics were comparable (**Fig. 4k,l**). These results show that neuronal activation drives a rapid rise in extracellular ATP that precedes adenosine accumulation, a sequence consistent with activity-dependent ATP release acting as a metabolic source of extracellular adenosine.

### Dual-color imaging of ATP and adenosine across sleep-wake cycles and movement

Having found that optogenetically evoked ATP release precedes adenosine accumulation in the BF, we asked whether the same sequence holds during spontaneous brain-state transitions. Simultaneous dual-color fiber photometry with ATP2h and rAdo1h in the BF showed that both signals track brain state, with ATP and adenosine levels significantly higher during wakefulness and REM sleep than during NREM sleep (**Fig. 5a**–**d**). As in the optogenetic experiments, ATP showed faster onset kinetics than adenosine during NREM-to-REM and REM-to-wake transitions, and faster decay kinetics during REM-to-wake transitions, with no significant kinetic differences during wake-to-NREM or NREM-to-wake transitions (**Fig. 5e,f**). Cross-correlation analysis showed that ATP2h reached its peak before rAdo1h at each transition (**Fig. 5g**), showing that spontaneous ATP release precedes adenosine accumulation in the BF during natural state changes.

**Fig. 5:**
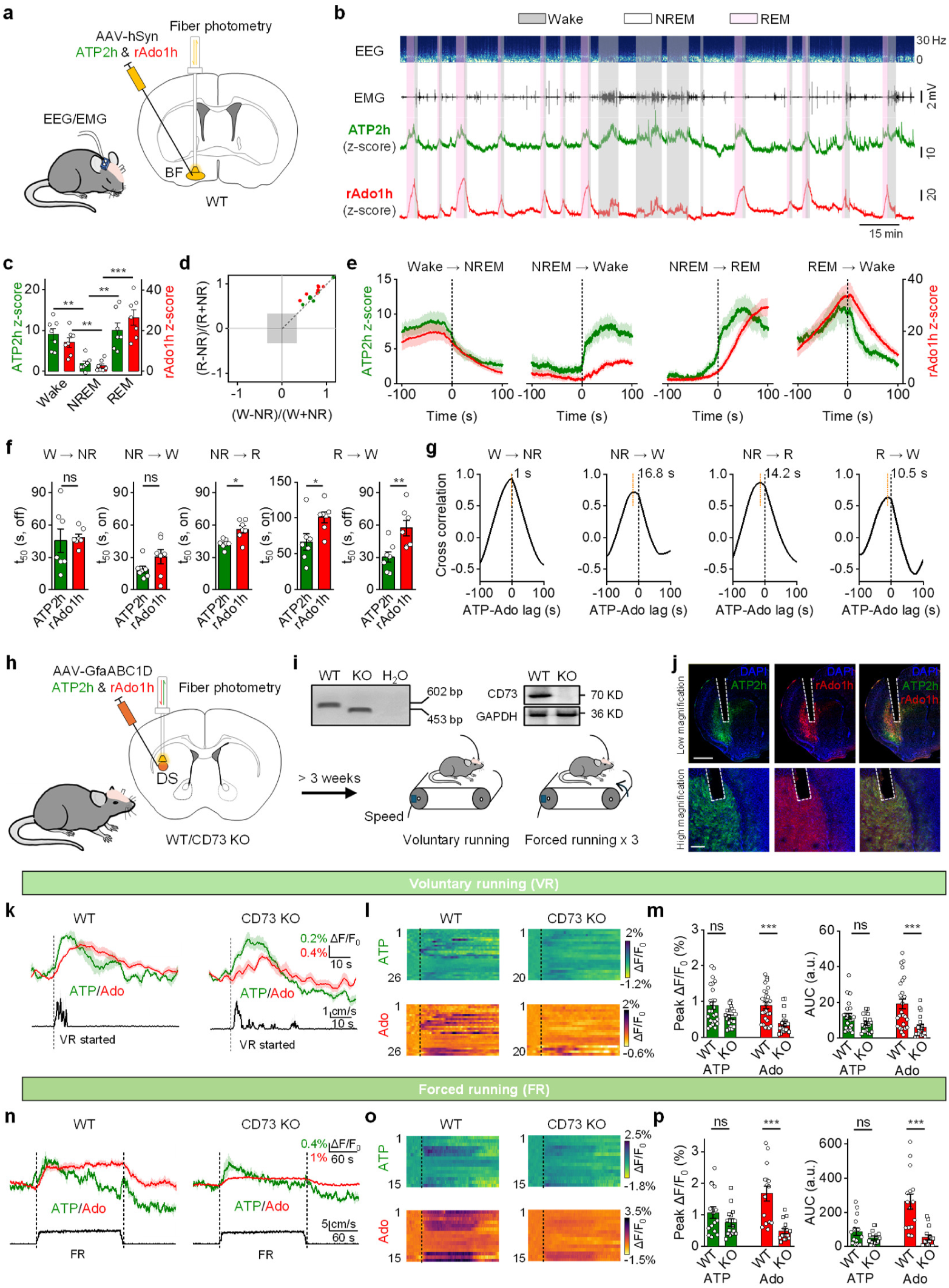
*In vivo* dual-color recording of ATP2h and rAdo1h during sleep-wake cycles and running. **a,** Schematic of dual-color fiber photometry recording of ATP2h (green) and rAdo1h (red) in the BF of freely moving mice, with EEG/EMG for sleep-state classification. **b**, Representative simultaneous EEG, EMG, and fluorescence traces for ATP2h and rAdo1h during spontaneous sleep-wake cycles. Shading: wakefulness (grey), REM sleep (pink), NREM sleep (unshaded). **c**, Summary of ATP2h and rAdo1h fluorescence (z-score) by vigilance state. *P* = 0.002 (ATP2h, wake vs. NREM), *P* = 0.001 (rAdo1h, wake vs. NREM), *P* = 0.001 (ATP2h, NREM vs. REM), *P* < 0.001 (rAdo1h, NREM vs. REM). Each symbol is one recording session; *n* = 7 sessions from 5 mice for c–g. **d**, Normalized signal modulation for each sensor across state comparisons (REM−NREM and Wake−NREM); grey box marks values under a twofold change. **e**, Time-aligned average fluorescence for ATP2h and rAdo1h centered on sleep-wake transitions (time 0, dashed line). **f**, Statistical comparison of the signal changes in e. *P* = 0.71 (W → NR), *P* = 0.32 (NR → W), *P* = 0.02 (NR → R), *P* = 0.04 for ton and *P* = 0.007 for toff (R → W). **g**, Cross-correlation of ATP2h and rAdo1h fluorescence aligned to the indicated transitions. Dashed black lines mark zero lag, orange lines mark peak cross-correlation lag, with peak-lag values shown above each plot. **h**, Schematic of dual-color fiber photometry recording of ATP2h (green) and rAdo1h (red) in the dorsal striatum (DS) of head-fixed mice during treadmill running. **i**, Validation of CD73 knockout (CD73 KO) at the genomic (left) and protein (right) levels. **j**, Representative low-magnification images (top) and separate high-magnification images (bottom) showing ATP2h (green) and rAdo1h (red) expression in the DS; nuclei were stained with DAPI (blue). Scale bar, 1 mm (top), 200 μm (bottom). **k**, Average fluorescence (top) and running traces (bottom) from a WT (left) and CD73 KO (right) mouse; dashed line marks onset of voluntary running. **l**, Pseudocolored ATP2h and rAdo1h responses to 26 voluntary running events in WT mice (left) and 20 events in CD73 KO mice (right). **m**, Summary of peak fluorescence change (left) and area under the curve (AUC, right) for each sensor during voluntary running, comparing WT and CD73 KO mice: *P* = 0.07 (ATP2h peak), *P* < 0.001 (rAdo1h peak), *P* = 0.90 (ATP2h AUC), *P* < 0.001 (rAdo1h AUC). *n* = 26 events from 5 WT mice and *n* = 20 events from 5 CD73 KO mice, respectively. **n**, Average fluorescence (top) and running traces (bottom) from a WT (left) and CD73 KO (right) mouse; dashed lines mark the duration of forced running. **o**, Pseudocolored ATP2h and rAdo1h responses to 15 forced running events in WT mice (left) and 15 events in CD73 KO mice (right). **p**, Summary of peak fluorescence change (left) and AUC (right) for each sensor during forced running, comparing WT and CD73 KO mice: *P* > 0.99 (ATP2h peak), *P* < 0.001 (rAdo1h peak), *P* > 0.99 (ATP2h AUC), *P* < 0.001 (rAdo1h AUC). *n* = 15 events from 5 WT mice and *n* = 15 events from 5 CD73 KO mice, respectively. Data are presented as mean ± s.e.m. Each data point represents one sample. Statistical significance was assessed using the Mann-Whitney *U* test for **c** and **f** and two-way ANOVA followed by Bonferroni’s multiple-comparisons test for **m** and **p**, listed *P* values are adjusted WT-vs.-CD73 KO comparisons within each sensor. Peak fluorescence change and AUC were analyzed separately. ns, not significant; \**P <* 0.05, \*\**P <* 0.01, \*\*\**P <* 0.001.

Adenosine also contributes to movement control in the dorsal striatum^35,36^. Using ATP2h and rAdo1h, we recorded striatal purinergic dynamics during voluntary and forced running (**Fig. 5h**–**j**). Consistent with earlier recordings using the green sensor Ado1.0^35^, rAdo1h tracked movement-associated adenosine release (**Fig. 5k**–**p**). To test whether CD73-dependent extracellular purine metabolism contributes to these adenosine signals, we examined mice lacking CD73, the ectonucleotidase that converts extracellular AMP to adenosine. In CD73 knockouts, rAdo1h responses were partially reduced during both voluntary and forced running, while ATP2h responses were unaffected (**Fig. 5k**–**p**). The A_2A_R-based rAdo1h sensor showed comparable JMF1907-induced responses in wild-type and CD73 knockout mice (**Extended Data Fig. 7**), arguing against gross genotype-dependent differences in sensor responsiveness and supporting the interpretation that the reduced running-associated adenosine signal reflects impaired CD73-dependent adenosine production.

Together, these experiments show that ATP and adenosine signals are temporally coupled *in vivo*. In the BF, ATP signals consistently preceded adenosine accumulation during optogenetic activation and sleep-wake transitions. In the dorsal striatum, CD73-dependent extracellular metabolism accounted for part of the movement-associated adenosine signal. These results support extracellular ATP-to-adenosine conversion as an important contributor to purinergic signaling *in vivo* and illustrate how the expanded GRAB sensor palette can resolve the spatiotemporal structure of purinergic signaling.

### ATP and adenosine dynamics in visual cortex during seizures

Purinergic signaling can both suppress and promote seizure activity^37^: Adenosine dampens hypersynchronous firing largely through A_1_ receptors^38^, while ATP can enhance excitability via P2X and P2Y receptors and also serves as the metabolic precursor that ectonucleotidases convert to adenosine^4,5^. The effects of purinergic signaling on seizures may depend on the relative timing of ATP release, adenosine production and adenosine clearance. Having observed temporally coupled ATP and adenosine signals during sleep-wake transitions and movement, we asked how these dynamics unfold during seizures.

We co-expressed the red ATP sensor rATP1h and the green adenosine sensor Ado2h in the visual cortex and used two-photon imaging to map ATP and adenosine dynamics at high spatiotemporal resolution before and after electrographic seizures (**Fig. 6a,b**). Both sensors showed large, seizure-locked fluorescence increases, with no response from the corresponding mutant sensors (**Fig. 6c**–**h**). The adenosine sensor exhibited slower rise and decay kinetics compared with the ATP sensor (**Fig. 6i,j**), but adenosine transients showed a distinctly prolonged late phase that persisted well past the point where the ATP signal had already decayed (**Fig. 6k**). This prolonged signal is consistent with sustained extracellular adenosine that remains present after ATP has returned to baseline, though the present data do not distinguish among the possible sources of that continued signal.

**Fig. 6:**
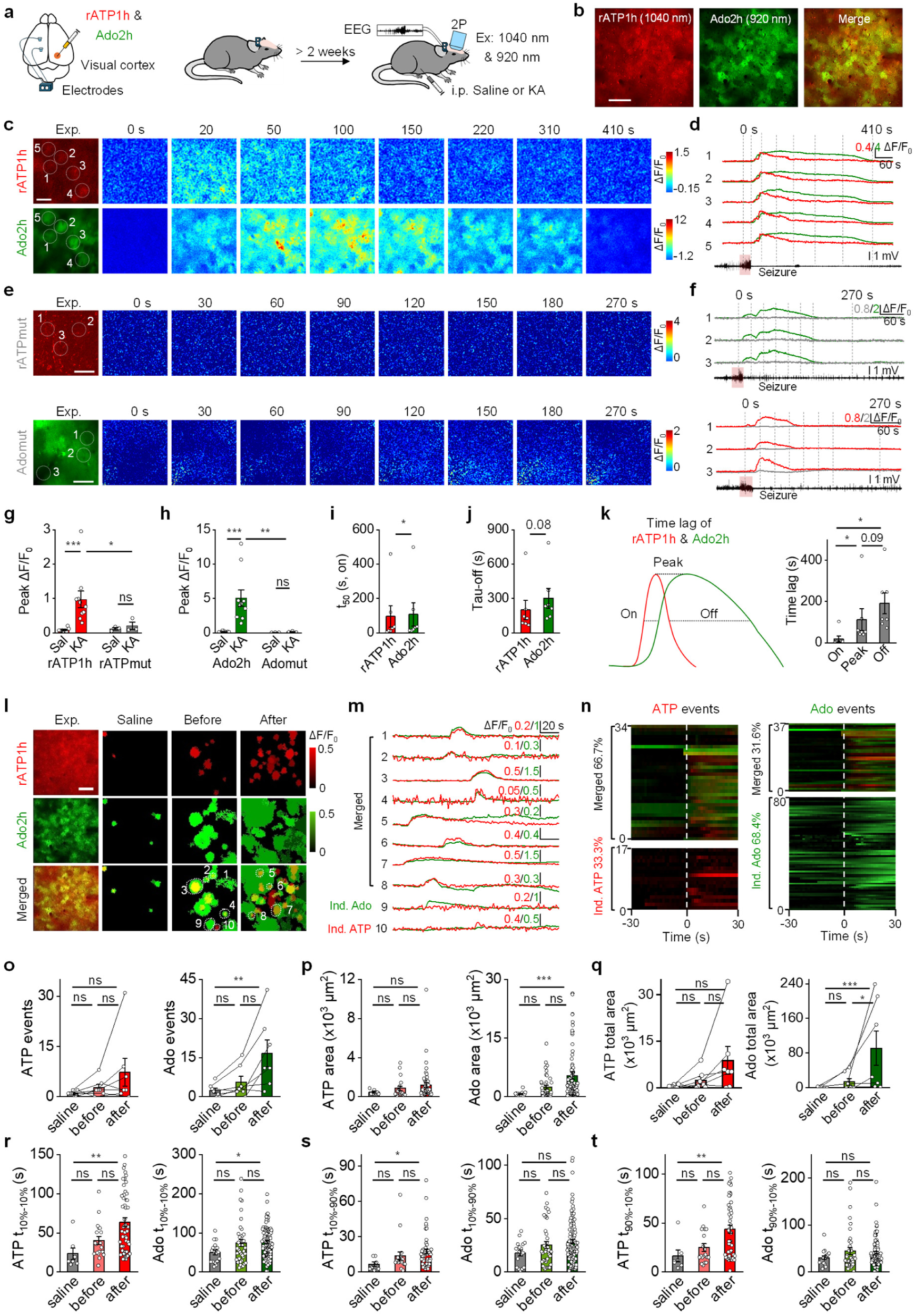
*In vivo* dual-color two-photon imaging of Ado2h and rATP1h in a mouse seizure model. **a,** Schematic of viral injection of AAV-GfaABC1D-Ado2h and AAV-GfaABC1D-rATP1h and head-fixed two-photon imaging in visual cortex (V1) during kainic acid (KA)-induced seizure. **b,** Representative *in vivo* two-photon image of V1 showing Ado2h and rATP1h fluorescence. Scale bar, 100 μm. **c**, Representative images of rATP1h (top) and Ado2h (bottom) expression and responses during seizure. Scale bar, 50 μm. **d**, Traces from the indicated ROIs (white circles) during seizure. **e,f,** As in **c,d**, but for mice injected with rATPmut virus (top) or Adomut virus (bottom). **g**, Peak fluorescence changes for rATP1h and rATPmut under saline and KA conditions. *n* = 10 and 3 mice for rATP1h and rATPmut, respectively. Two-way mixed-design ANOVA: sensor effect, *P* = 0.155; treatment effect, *P* = 0.00256; sensor × treatment interaction, *P* = 0.0889. Bonferroni-adjusted planned comparisons: *P* = 0.0107 (rATP1h, saline vs. KA), *P* > 0.999 (rATPmut, saline vs. KA) and *P* = 0.0426 (rATP1h vs. rATPmut under KA). **h**, Peak fluorescence changes for Ado2h and Adomut under saline and KA conditions. *n* = 10 and 3 mice for Ado2h and Adomut, respectively. Two-way mixed-design ANOVA: sensor effect, *P* = 0.0481; treatment effect, *P* = 0.00209; sensor × treatment interaction, *P* = 0.0559. Bonferroni-adjusted planned comparisons: *P* = 0.00801 (Ado2h, saline vs. KA), *P* = 0.407 (Adomut, saline vs. KA) and *P* = 0.00793 (Ado2h vs. Adomut under KA). **i**, Rise t_50_ of Ado2h and rATP1h during seizures. *n* = 7 mice; *P* = 0.0611. **j**, Decay time constant of Ado2h and rATP1h during seizures. *n* = 7 mice; *P* = 0.0284. **k**, Schematic (left) and quantification (right) of the temporal lag between the Ado2h and rATP1h signals at signal onset, peak and offset. *n* = 7 mice. Friedman test, *P* = 0.00215; Bonferroni-adjusted pairwise Wilcoxon signed-rank tests: *P* = 0.0469 (onset vs. peak), *P* = 0.0469 (onset vs. offset) and *P* = 0.0938 (peak vs. offset). **l**, Representative *in vivo* two-photon images of rATP1h (red) and Ado2h (green) under saline conditions and before/after seizure under KA. Scale bar, 100 μm. **m**, Representative ΔF/F_0_ traces of rATP1h (red) and Ado2h (green) from individual ROIs (ROI1–10, white circles in **l**) before and after seizure. **n**, Time-aligned heatmaps of ATP events (left) and Ado events (right); percentages indicate the proportion of merged events relative to total events, aligned to event onset (t = 0 s). **o**, Numbers of ATP and adenosine events under saline, before-seizure and after-seizure conditions. *n* = 7 mice. ATP: Friedman test, *P* = 0.223. Adenosine: Friedman test, *P* = 0.00726; Dunn’s multiple-comparisons test, *P* = 0.544 (saline vs. before), *P* = 0.00635 (saline vs. after) and *P* = 0.247 (before vs. after). **p**, Areas of individual ATP and adenosine events under the same conditions. ATP: Kruskal-Wallis test, *P* = 0.120. Adenosine: Kruskal-Wallis test, *P* = 0.00716; Dunn’s multiple-comparisons test, *P* = 0.240 (saline vs. before), *P* = 0.00811 (saline vs. after) and *P* = 0.434 (before vs. after). **q**, Total areas of ATP and adenosine events under the same conditions. *n* = 7 mice. ATP: Friedman test, *P* = 0.102. Adenosine: Friedman test, *P* = 0.00373; Dunn’s multiple-comparisons test, *P* > 0.999 (saline vs. before), *P* = 0.00635 (saline vs. after) and *P* = 0.0334 (before vs. after). **r**, Event duration (t10–10%) for ATP and Ado across the same conditions. ATP: Kruskal-Wallis test, *P* = 0.00219; Dunn’s multiple-comparisons test, *P* = 0.258 (saline vs. before), *P* = 0.00414 (saline vs. after) and *P* = 0.145 (before vs. after). Adenosine: Kruskal-Wallis test, *P* = 0.0124; Dunn’s multiple-comparisons test, *P* = 0.295 (saline vs. before), *P* = 0.0140 (saline vs. after) and *P* = 0.510 (before vs. after). **s**, Rising-phase duration (t10–90%) for ATP and Ado across the same conditions. ATP: Kruskal-Wallis test, *P* = 0.0122; Dunn’s multiple-comparisons test, *P* = 0.510 (saline vs. before), *P* = 0.0225 (saline vs. after) and *P* = 0.239 (before vs. after). Adenosine: Kruskal-Wallis test, *P* = 0.106. **t**, Decay-phase duration (t90–10%) for ATP and Ado across the same conditions. ATP: Kruskal-Wallis test, *P* = 0.00188; Dunn’s multiple-comparisons test, *P* = 0.521 (saline vs. before), *P* = 0.00694 (saline vs. after) and *P* = 0.0593 (before vs. after). Adenosine: Kruskal-Wallis test, *P* = 0.345. For **p** and **r**–**t**, *n* = 7, 19 and 51 ATP events and *n* = 17, 39 and 117 adenosine events under saline, before-seizure and after-seizure conditions, respectively; events were pooled from 7 mice. Data are presented as mean ± s.e.m. Each point represents one mouse in **g–k**, **o** and **q** and one event in **p** and **r–t**. All tests were two-sided. Two-way mixed-design ANOVA with Bonferroni-corrected planned comparisons (**g**,**h**); Two-sided paired *t*-test (**i**,**j**); Friedman test followed by the indicated corrected post hoc comparisons (**k**,**o**,**q**); Kruskal-Wallis test followed, where appropriate, by Dunn’s multiple-comparisons test (**p**,**r–t**); ns, not significant, \**P <* 0.05, \*\**P <* 0.01, \*\*\**P <* 0.001.

To exclude the possibility that the prolonged adenosine signal reflected the specific sensor pair, we repeated the experiment using ATP2h and rAdo1h (**Extended Data Fig. 8a,b and Supplementary Video 1**). Both sensors showed seizure-associated responses, and rAdo1h again displayed a prolonged signal after the ATP2h signal had declined (**Extended Data Fig. 8c,d**). The corresponding mutant sensors showed no response (**Extended Data Fig. 8e–h**). ATP2h and rAdo1h showed similar rise kinetics, whereas rAdo1h decayed more slowly (**Extended Data Fig. 8i–k**), supporting the sustained adenosine phase with an alternative sensor pair.

The spatial resolution afforded by the genetically encoded sensors enabled analysis of spontaneous purinergic events at individual active sites. After seizures, ATP and adenosine events became more frequent, covered larger areas and showed slower kinetics (**Fig. 6o**–**t**). Approximately 67% of ATP events spatially overlapped with adenosine events, whereas approximately 32% of adenosine events overlapped with ATP events (**Fig. 6n**). This asymmetry indicates that ATP events often spatially overlap with adenosine events, whereas many adenosine events lack a detectable coincident ATP event. Such non-overlapping adenosine events could reflect ATP-independent adenosine release, ATP signals below the detection threshold or spatial separation between ATP release and adenosine production.

### Dual-color imaging of extracellular and intracellular adenosine reveals compartment-specific dynamics during seizures

Adenosine can be released directly from neurons rather than arising solely from extracellular ATP breakdown^19,39^. To test whether intracellular and extracellular adenosine dynamics are coordinated, we co-expressed the recently developed intracellular sensor HypnoS^40^ with the extracellular sensor rAdo1h in cultured neurons. Electrical stimulation drove concurrent increases in both signals that scaled with stimulation intensity, and the two were closely correlated, with the intracellular signal rising ahead of the extracellular one (**Extended Data Fig. 9a–h**). These culture experiments established the compatibility of the two sensors for paired recordings but did not determine the direction of adenosine transfer between compartments. We next examined their relative dynamics *in vivo*, where compartment-specific patterns are more relevant to seizure-associated pathology.

Extracellular adenosine levels reflect a combination of production, transport, and metabolism^41–47^, and it remains unclear whether neuronal, astrocytic and extracellular adenosine follow the same temporal profiles during pathological activity. To address this question, we co-expressed rAdo1h with HypnoS targeted to either neurons or astrocytes and monitored extracellular and intracellular adenosine signals during seizures. Seizure-associated increases in extracellular rAdo1h fluorescence coincided with HypnoS responses in both neurons and astrocytes (**Fig. 7a**–**j**). In seizure-aligned traces, extracellular rAdo1h fluorescence increased before intracellular HypnoS fluorescence in both neuronal and astrocytic recordings **(Fig. 7d,h**). The extracellular signal also decayed more slowly than either intracellular signal (**Fig. 7l**), and peak-time analysis revealed distinct temporal offsets between extracellular adenosine and neuronal or astrocytic intracellular adenosine (**Fig. 7m**). These results show that extracellular, neuronal intracellular and astrocytic intracellular adenosine follow distinct temporal profiles during seizures, consistent with compartment-specific handling of adenosine rather than a single shared kinetic process.

**Fig. 7:**
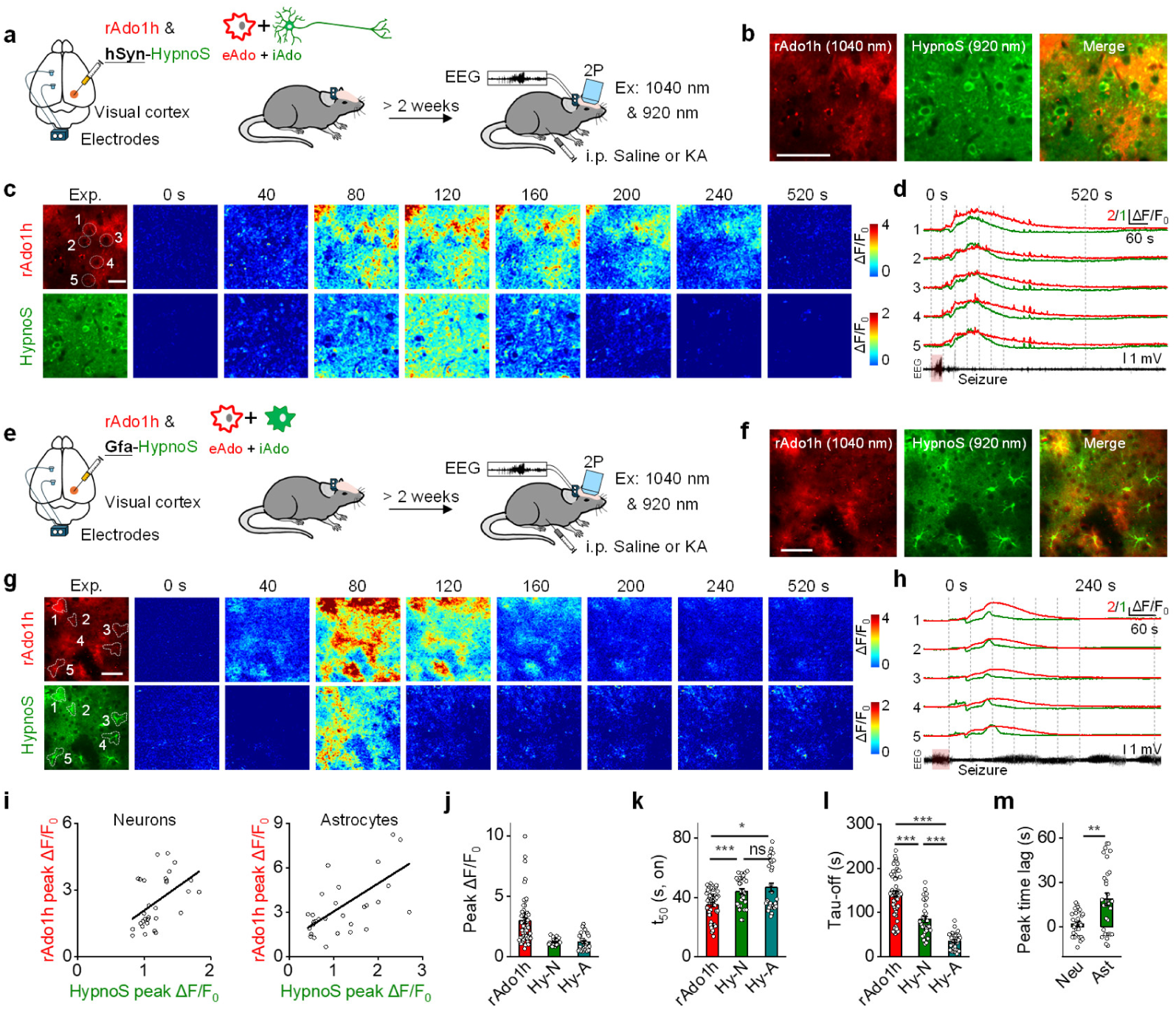
*In vivo* dual-color two-photon imaging of extracellular and intracellular adenosine in a mouse seizure model. **a,** Schematic of viral injection of rAdo1h and neuron-specific HypnoS with head-fixed two-photon imaging in V1 during KA-induced seizure. **b**, Representative *in vivo* two-photon image of V1 showing rAdo1h and HypnoS fluorescence. Scale bar, 50 μm. **c**, Representative images of rAdo1h (top) and HypnoS (bottom) expression and responses during seizure. Scale bar, 50 μm. **d**, Traces from the indicated ROIs (white circles) during seizure. **e**–**h**, As in **a–d**, but with HypnoS expressed in astrocytes. **i**, Correlation between peak fluorescence changes of rAdo1h and HypnoS in neurons (left) and astrocytes (right). Neuronal HypnoS: Pearson’s *r*(28) = 0.486, two-sided *P* = 0.00652, 95% CI [0.152, 0.720]. Astrocytic HypnoS: Pearson’s *r*(28) = 0.590, two-sided *P <* 0.001, 95% CI [0.292, 0.784]. *n* = 30 ROIs from 3 mice per group. **j**, Peak fluorescence changes of rAdo1h, neuronal HypnoS and astrocytic HypnoS during seizures. Kruskal-Wallis test, *P* < 0.001; Dunn’s multiple-comparisons test, *P* < 0.001 (rAdo1h vs. Hy-N), *P* < 0.001 (rAdo1h vs. Hy-A) and *P* > 0.999 (Hy-N vs. Hy-A). **k**, Rise t_50_ of rAdo1h, neuronal HypnoS and astrocytic HypnoS during seizures. Kruskal-Wallis test, *P* = 0.00187; Dunn’s multiple-comparisons test, *P* = 0.00471 (rAdo1h vs. Hy-N), *P* = 0.0309 (rAdo1h vs. Hy-A) and *P* > 0.999 (Hy-N vs. Hy-A). **l**, Decay time constants of rAdo1h, neuronal HypnoS and astrocytic HypnoS during seizures. Kruskal-Wallis test, *P* < 0.001; Dunn’s multiple-comparisons test, *P* < 0.001 for all three pairwise comparisons. **m**, Peak-time lag between the rAdo1h and HypnoS signals in neuronal and astrocytic recordings (lag = t_peak, rAdo1h_ − t_peak, HypnoS_). *n* = 30 ROIs per group; *P* = 0.00281. For **j**–**l**, *n* = 60, 30 and 30 ROIs for rAdo1h, neuronal HypnoS and astrocytic HypnoS, respectively, pooled from 3 mice per experimental group. Hy-N, neuronal HypnoS; Hy-A, astrocytic HypnoS. Data are presented as mean ± s.e.m. Each point represents one ROI. All tests were two-sided. Pearson correlation (**i**); Kruskal-Wallis test followed by Dunn’s multiple-comparisons test (**j**–**l**); Mann-Whitney *U* test (**m**); ns, not significant, \**P <* 0.05, \*\**P <* 0.01, \*\*\**P <* 0.001.

## Discussion

Genetically encoded fluorescent sensors have reshaped how neuromodulatory signaling is studied, making it possible to image neurochemical activity in specific cell types with fine spatial and temporal resolution in behaving animals^16,17,48^. The work presented here extends the GRAB platform with a matched set of ATP and adenosine sensors built specifically for multiplexed imaging of purinergic signaling *in vivo*. Systematic engineering across receptor scaffolds, membrane trafficking, and fluorophore coupling produced green sensors and spectrally separated red counterparts with brighter output, quicker kinetics, tight ligand specificity, and little engagement of native GPCR pathways (**Fige. 1 and 2**). Taken together, these properties allow ATP and adenosine to be tracked in parallel, with minimal disturbance to the very signaling events under observation^16^.

What sets this toolkit apart is the ability to follow two metabolically intertwined purines side by side^9^. Existing genetically encoded options^18,20,41^ can report ATP or adenosine, but not both at once; they occupy the same spectral window and lack the temporal resolution needed to separate the two signals in time. Splitting the GRAB palette into paired green and red variants solves this, yielding a dual-color platform compatible with fiber photometry and two-photon microscopy alike. This makes it possible to watch ATP release and adenosine accumulation unfold together in the same circuit, giving a direct window onto how extracellular purinergic signaling is organized in time.

Applying this approach, we found that ATP transients reliably preceded extracellular adenosine accumulation during neuronal activation (**Fig. 4f**–**l**), and again across sleep-wake transitions (**Fig. 5**). In the dorsal striatum, CD73 knockout partially reduced adenosine responses during voluntary and forced running without affecting ATP responses (**Fig. 5k**–**p**), indicating that CD73-dependent hydrolysis accounts for only part of the movement-evoked adenosine signal and that other sources, potentially including direct adenosine release, contribute as well. We did not perform equivalent CD73 or CD39 perturbations in the BF sleep-wake recordings, CA1 optogenetic experiments or visual-cortex seizure experiments. Thus, in these settings, the temporal relationship between ATP and adenosine should be interpreted as consistent with, rather than evidence for, sequential enzymatic conversion. This distinction accords with earlier work describing sequential ATP breakdown through the CD39-CD73 pathway alongside equilibrative nucleoside transporter (ENT)-mediated efflux of intracellular adenosine from neurons^3,5,9,18,19,39,41–43^. This toolkit enables simultaneous, high-resolution measurement of ATP and adenosine signals in intact circuits. Establishing ATP-to-adenosine conversion in each setting will require CD73/CD39 perturbations in the corresponding preparation.

The expanded sensor palette also lets purinergic signaling be examined across spatial scales within a single study. Fiber photometry captured fast extracellular ATP and adenosine dynamics during ordinary behavior, while two-photon imaging resolved the heterogeneity of purinergic release at individual sites during seizures (**Fig. 6**). Pairing the extracellular GRAB sensors with the intracellular adenosine sensor HypnoS^40^ extended this further, allowing extracellular adenosine to be measured simultaneously with either neuronal or astrocytic intracellular adenosine **(Fig. 7)**. In seizure-aligned recordings, extracellular rAdo1h fluorescence increased before intracellular HypnoS fluorescence in both neuronal and astrocytic recordings, and the extracellular signal decayed more slowly than either intracellular signal (**Fig. 7d,h,l**). Peak-time analysis also revealed distinct temporal offsets between extracellular adenosine and neuronal or astrocytic intracellular adenosine (**Fig. 7m**). These findings reveal compartment-specific temporal profiles during seizures, but do not identify the cellular source of extracellular adenosine or the direction of adenosine transfer between compartments. More broadly, these experiments show how spectrally matched genetically encoded sensors can be combined across imaging modalities and cellular compartments to build a multiplexed view of metabolite dynamics spanning molecules, cells and circuits^49^.

Beyond validating the sensors, these experiments point to several features of purinergic organization worth further dissection. We observed rapid adenosine transients during acute stress (**Fig. 3l**–**r**); a recurring sequence in which ATP elevations preceded adenosine accumulation across neuronal activation and sleep-wake transitions (**Fige. 4 and 5**); and an extracellular adenosine signal that, during seizures, persisted substantially longer than ATP (**Fig. 6**). The ATP-before-adenosine sequence observed during neuronal activation and sleep-wake transitions is consistent with extracellular ATP-to-adenosine conversion via ectonucleotidase pathways^44,45^, though direct, ATP-independent adenosine release may also contribute. The compartment-resolved seizure data add a further layer to this picture: extracellular, neuronal intracellular and astrocytic intracellular adenosine followed distinct rise, peak and decay profiles during seizures (**Fig. 7**). These differences are consistent with compartment-specific adenosine handling, but do not establish whether either intracellular compartment supplies extracellular adenosine. These temporal differences should be interpreted in the context of established mechanisms of adenosine production, transport and metabolism in neural tissue^43,46,47^. Together, these findings show that simultaneous recording of multiple purines can resolve temporal relationships and compartment-specific dynamics that single-signal recordings would miss.

The current red sensors already support multiplexed imaging, though their responses remain smaller in magnitude than those of the optimized green variants (**Fig. 1i,j**). Further engineering of red and far-red variants^50,51^ could improve sensitivity and create room to pair purine imaging with additional signaling modalities. The GRAB platform’s modular design also leaves room to extend multiplexed imaging to other neuromodulators and metabolites; as the spectral range of genetically encoded sensors continues to expand, tracking several extracellular and intracellular pathways at once within the same preparation should become increasingly practical.

In summary, we present an expanded GRAB sensor toolkit for multiplexed imaging of purinergic signaling *in vivo*. By combining improved sensor performance with spectral diversity and cross-modality compatibility, this toolkit supports direct measurement of ATP and adenosine dynamics at high spatiotemporal resolution in behaving animals and generates testable predictions about extracellular ATP-to-adenosine conversion and compartment-specific adenosine handling during seizures. We expect this toolkit to serve as a general resource for studying how purinergic signaling shapes neural circuit function in health and disease.

### Methods Molecular biology

Coding sequences for human adenosine A_2A_ receptor (A_2A_R), chicken P2Y_1_ receptor (P2Y_1_R), and zebra finch P2Y_1_R were obtained as follows: human A_2A_R was cloned from HEK293T cell cDNA, chicken P2Y_1_R was cloned from chicken genomic DNA extracted from skeletal muscle tissue, and the zebra finch P2Y_1_R coding sequence was synthesized based on the UniProt sequence H0ZM60. Plasmids were constructed using an in-house Gibson assembly protocol, and all sequences were verified by Sanger sequencing. For screening and characterization in HEK293T cells, the fluorescent sensors were subcloned into the pDisplay vector (Invitrogen). Site-directed mutagenesis was performed using primers containing randomized NNK codons (48 codons in total, encoding all 20 amino acids; Beijing Xianghong Biological Technology) or defined codons. For characterization in cultured neurons, sensor genes were cloned into pAAV vectors under the control of the human synapsin (hSyn) promoter or the GFAP-derived GfaABC1D promoter. For luciferase complementation assays, GPCR-SmBit or sensor-SmBit constructs were generated based on β_2A_R-SmBit^29^. The LgBit-miniGs and LgBit-miniGsq plasmids were constructed based on the pBiT1.1 vector (Promega). The LgBit-miniGs and LgBit-miniGsq plasmids were constructed in the pBiT1.1 vector (Promega), with miniGs or miniGsq fused downstream of LgBit through a GSSGGGGSGGGGSS flexible linker. For Tango assays, the corresponding GPCR and sensor constructs were generated in the pTango backbone from the PRESTO-Tango kit. For sensor-Tango constructs, the N-terminal IgK or LucyRho signal peptide was removed before cloning because the pTango vector already contains an N-terminal hemagglutinin signal peptide.

### Sensor library construction and screening

Sensor libraries for insertion-site screening were generated by combinatorial Gibson assembly in pDisplay-based screening vectors. For cpmApple-based sensors, an IRES-EGFP-CAAX cassette was included as a membrane marker for downstream analysis, whereas for cpEGFP-based sensors, an IRES-mCherry-CAAX cassette was used. For each candidate N-terminal insertion site, a site-specific reverse primer was designed to amplify the upstream GPCR fragment and append a common overlap sequence homologous to the 5′ end of cpEGFP or cpmApple. Similarly, for each candidate C-terminal insertion site in TM6, a site-specific forward primer was designed to amplify the downstream GPCR fragment and append a common overlap sequence homologous to the 3′ end of cpEGFP or cpmApple. The upstream GPCR fragment, downstream GPCR fragment, fluorescent protein module, and linearized vector backbone was assembled by

Gibson assembly and transformed into in-house-prepared DH5α competent cells. For insertion-site screening libraries, transformation scale was adjusted to obtain approximately 2- to 3-fold coverage of all possible N- and C-terminal insertion-site combinations. For site-directed mutagenesis libraries, 47 colonies were picked for each screen.

Individual bacterial colonies were picked into 96-well deep-well plates containing 1.2 ml LB medium supplemented with ampicillin and grown overnight. Each plasmid variant was prepared individually in 96-well format. Plasmid extraction was performed using in-house-prepared miniprep buffers together with 96-well plasmid purification columns from the TIANprep N96 Plasmid Kit (Tiangen Biotech). Purified plasmids were eluted and used directly for cell transfection and sensor screening.

Detailed sensor-specific engineering procedures and optimization trajectories are provided in Supplementary information.

### Cell culture

HEK293T cells were cultured at 37 °C in humidified air containing 5% CO_2_ in DMEM (Procell) supplemented with 10% (vol/vol) fetal bovine serum (FBS; Vazyme, F101) and 1% penicillin-streptomycin (Procell). When cells reached approximately 70% confluence, they were transfected with PEI using 1 µg plasmid DNA and 3 µg PEI per well in 24-well plates. Imaging was performed 24–48 h after transfection.

Primary rat cortical neurons were prepared from postnatal day 0 (P0) Sprague-Dawley rats. Briefly, cortical tissue was dissected, enzymatically digested with 0.25% trypsin-EDTA (Biological Industries), mechanically dissociated, and plated onto poly-D-lysine-coated 12-mm glass coverslips (Sigma-Aldrich) in 24-well plates. Neurons were maintained at 37 °C in humidified air containing 5% CO_2_ in Neurobasal medium (Thermo) supplemented with 2% B-27 (Thermo), 1% GlutaMAX (Thermo), and 1% penicillin-streptomycin. Cultured neurons were transduced with AAVs encoding the indicated sensors at 3–5 days *in vitro* (DIV3–5) and imaged at DIV11–14.

### Fluorescence imaging in cultured cells

Fluorescence imaging of cultured cells was performed after replacing the culture medium with Tyrode’s solution containing (in mM): 150 NaCl, 4 KCl, 2 MgCl_2_, 10 HEPES, and 10 glucose (pH 7.35–7.45 adjusted with NaOH). Cells were imaged in a custom-made chamber using either an inverted Ti-E A1 confocal microscope (Nikon) controlled by NIS-Elements 4.51.00 software (Nikon) or an Opera Phenix high-content screening system (PerkinElmer) with Harmony 4.9 software (PerkinElmer). The confocal system was equipped with ×10/0.45 NA, ×20/0.75 NA, and ×40/1.35 NA oil-immersion objectives, 488-nm and 561-nm lasers, and emission filters of 525/50 nm (green) and 595/50 nm (red). During imaging, compounds including SCH-58261 (MedChemExpress, MCE; A_2A_R antagonist), MRS-2500 (MCE; P2Y_1_R antagonist), adenosine triphosphate (ATP; MCE), adenosine diphosphate (ADP; MCE), adenosine monophosphate (AMP; MCE), adenosine (Ado; MCE), adenine (Ade; MCE), inosine (Ino; MCE), uridine triphosphate (UTP; MCE), uridine diphosphate (UDP; MCE), UDP-glucose (UDP-G; MCE), uridine monophosphate (UMP; MCE), guanosine triphosphate (GTP; MCE), glutamate (Sigma-Aldrich), γ-aminobutyric acid (GABA; Tocris), glycine (Sigma-Aldrich), dopamine (Sigma-Aldrich), norepinephrine (NE; Tocris), serotonin (5-HT; MCE), and acetylcholine (ACh; Solarbio) were applied either by bath application or using a custom perfusion system at the indicated concentrations. Fluorescence changes were quantified as ΔF/F_0_ = (F – F_0_)/F_0_, with F_0_ defined as the mean baseline fluorescence over 0–1 min prior to drug application. The Opera Phenix system was equipped with ×20/1.0 NA, ×40/0.6 NA, and ×40/1.15 NA water-immersion objectives, with 488-nm and 561-nm excitation lasers; green fluorescence was collected using a 525/50 nm emission filter and red fluorescence using a 600/30 nm emission filter.

### Spectral measurements

Plasmids encoding GRAB_Ado_ and GRAB_ATP_ sensors were transfected into HEK293T cells seeded in six-well plates. Twenty-four to thirty hours after transfection, cells were collected, resuspended in assay buffer, and transferred to 384-well plates. Basal fluorescence was measured in assay buffer supplemented with adenosine deaminase (ADA; Shanghai Yuanye) for Ado sensors or apyrase grades VI and VII (Sigma-Aldrich) for ATP sensors to remove ambient Ado or ATP, respectively. For maximum fluorescence measurements, ADA or apyrase was not added; instead, 100 μM Ado or ATP was directly added to the corresponding sensor preparation. Excitation and emission spectra were acquired using a Safire2 multimode plate reader (Tecan) with 5-nm wavelength increments. Fluorescence signals from non-transfected cells were measured in parallel and subtracted as background. All measurements were performed at room temperature under identical instrument settings.

### Photoactivation measurements

Cells expressing the indicated fluorescent sensors were cultured on coverslips. Before imaging, culture medium was replaced with 4K buffer. Measurements were performed without exogenously added sensor ligands. Bursts of 488-nm laser light (1-s duration, 200 μW, approximately 0.4 W/cm^2^) were delivered to induce photoactivation. Fluorescence imaging was paused during blue-light stimulation and resumed immediately afterward. Cellular regions of interest were defined in ImageJ. Background fluorescence was automatically estimated from dark, non-cellular pixels outside the cellular ROIs and subtracted from each frame. To correct cumulative baseline drift during repeated stimulation, the baseline of each trial was estimated as the median fluorescence during the final five pre-stimulation frames. For each cell, the difference between this baseline and the first-trial baseline was subtracted as a constant offset from every frame of that trial. Thus, correction aligned trial baselines without fitting or altering within-trial fluorescence kinetics. For each trial, F_0_ was calculated as the mean corrected fluorescence across all pre-stimulation frames, and fluorescence responses were expressed as ΔF/F_0_ = (F – F_0_)/F_0_. Photoactivation amplitude was defined as the maximum single-frame ΔF/F_0_ observed within 3 s after the end of blue-light stimulation. Peak responses from repeated trials were averaged to obtain one photoactivation value per cell.

### Luciferase complementation assay

HEK293T cells were co-transfected with the indicated GRAB_Ado_ and GRAB_ATP_ sensors or WT receptors together with the corresponding LgBiT-mG construct at 50–60% confluence. Approximately 24–36 h after transfection, cells were dissociated using a cell scraper, resuspended in PBS, and plated into 96-well plates. NECA or ADPβS at concentrations ranging from 0.1 nM to 1 mM, together with 5 μM furimazine (Nano-Glo Luciferase Assay, Promega), were bath-applied to the cells. After a 10-min incubation at room temperature in the dark, luminescence signals were measured using a VICTOR X5 multi-label plate reader (PerkinElmer). Dose-response curves were generated to evaluate sensor sensitivity and dynamic range.

### Membrane preparation and stopped-flow kinetic measurements

For kinetic measurements, GRAB sensor coding sequences were subcloned into a pPiggyBac-derived vector containing a puromycin-resistance cassette. HEK293T cells cultured in DMEM supplemented with 10% FBS and 1% penicillin-streptomycin were cotransfected in 10-cm dishes with 10 µg total DNA comprising the sensor vector and hyPBase plasmid at a 20:1 mass ratio. Transfection was performed using 40-kDa PEI at a PEI: DNA mass ratio of 3:1. Cells were selected with 2–4 µg/ml puromycin for 1 week to generate polyclonal stable sensor-expressing cell lines.

For each membrane preparation, two 15-cm dishes of stable cells at approximately 90% confluence were collected. Membranes were prepared using a protocol adapted from Culhane et al^52^. Cells were washed with PBS and resuspended in hypotonic buffer containing 10 mM HEPES and 50 mM EDTA, pH 7.4, supplemented with 1× Roche protease inhibitor cocktail. After incubation for 30 min, cells were disrupted using a chilled Dounce homogenizer. Cell debris and nuclei were removed by centrifugation at 1,000 g for 5 min at 4 °C, and membranes were pelleted from the supernatant by ultracentrifugation at 135,000 g for 25 min at 4 °C. DTT, PMSF, aprotinin, and leupeptin used in the original protocol were omitted and replaced with 1× Roche protease inhibitor cocktail.

Membrane pellets were resuspended in membrane buffer containing 20 mM HEPES, 5 mM KCl, 145 mM NaCl, 2 mM CaCl_2_, and 1 mM MgCl_2_, pH 7.4. To remove residual ambient ATP, membranes from the high-affinity ATP2h and rATP1h sensor lines were washed at least three times. During each wash, membrane pellets were resuspended in 1–2 ml membrane buffer and re-pelleted at 135,000 g. Before measurement, membrane preparations were resuspended in 15–20 ml 4K buffer containing 150 mM NaCl, 4 mM KCl, 2 mM MgCl_2_, 10 mM HEPES, and 10 mM glucose, adjusted to pH 7.35–7.45.

Fluorescence kinetics were measured at 25 °C using an Applied Photophysics SX20 stopped-flow spectrometer with a 1:1 mixing ratio. Green fluorescent sensors were excited at 490 nm, and fluorescence was collected through a 515/30-nm band-pass filter. Red fluorescent sensors were excited at 550 nm, and fluorescence was collected through a 590/50-nm band-pass filter.

For association measurements, membrane suspensions in 4K buffer were mixed with an equal volume of ligand prepared at twice the desired final concentration in the same buffer. Acquisition duration was adjusted empirically for each sensor so that fluorescence was recorded until the response reached an apparent plateau. Each ligand concentration or dissociation condition was measured at least three times.

Kinetic traces were analyzed in Origin. Association traces were fitted using the ExpGro1 model,

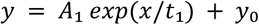

### Animals

All procedures for animal surgery and experimentation were performed in accordance with and approved by the laboratory animal care and use committees of the Institute of Genetics and Developmental Biology, Chinese Academy of Sciences (AP2024018). Hippocampal neurons were prepared from both male and female postnatal day 0 (P0) Sprague-Dawley rats (Charles River). Experiments also utilized P48–P90 wild-type C57BL/6N and C57BL/6J mice (Charles River), CaMKIIα-Cre mice (The Jackson Laboratory; JAX:016963), VGLUT2-Cre mice (The Jackson Laboratory; JAX:005359) and CD73 KO (The Jackson Laboratory; JAX:018986) mice. Animals were housed under a standard 12-h light/dark cycle at 22–25 °C with 40–60% humidity, with food and water available ad libitum. Animals were randomly assigned to experimental groups.

### Western blotting

Brain tissues were collected from wild-type and CD73 knockout (CD73 KO) mice and homogenized in RIPA buffer supplemented with protease inhibitors. Equal amounts of protein were separated by SDS-PAGE and transferred onto PVDF membranes. Membranes were blocked with 5% non-fat milk and incubated with primary antibodies against CD73 (ABclonal, A2029SP; 1:1,000) and GAPDH (loading control, Abmart, M20006; 1:10,000), followed by incubation with HRP-conjugated secondary antibodies (Abmart, M21001, M21002; 1:10,000). Protein bands were visualized using an enhanced chemiluminescence detection system.

### Histology

Mice were anesthetized and transcardially perfused with phosphate-buffered saline (PBS), followed by 4% paraformaldehyde (PFA) in PBS. The brains were post-fixed in 4% PFA at 4 °C overnight and then transferred to 30% sucrose/PBS at 4 °C for 48–72 h. Brains were sectioned on a freezing microtome (Leica, CM1950) and coronal sections (30 μm) were collected. To confirm the sensors’ expression at the fiber implantation site, brain sections were mounted with Antifade Mounting Medium (ACMEC, Lot. No. A53335E1E) on coverslips and directly imaged. To visualize DIO-ChrimsonR-HaloTag expression, mice received a retro-orbital injection of 200 nmol SiR650 24 h before perfusion to label HaloTag-expressing neurons for fluorescence imaging.

### AAV vectors

Recombinant AAV vectors were packaged by WZ Biosciences. Sensor and control sequences were expressed under the hSyn or GfaABC1D promoter and packaged in AAV2/9 or AAV2/5 capsids, respectively. The recombinant AAVs included Ado1.0, Ado2h, Ado2m, rAdo1h, Adomut, rAdomut, ATP1.0, ATP2h, ATP2l, rATP1h, ATPmut, rATPmut, DIO-GCaMP6s, DIO-ChrimsonR-HaloTag, and HypnoS. Viral titers ranged from 1.5 × 10^12^ to 9.8 × 10^13^ vg ml^−1^. Recombinant AAVs were used either for transduction of cultured neurons or for stereotaxic injection into specific brain regions for *in vivo* experiments.

### Stereotaxic AAV delivery and optical fiber implantation

Adult mice were anesthetized with isoflurane (2.5% for induction and 0.5–1% for maintenance) and secured in a stereotaxic frame (RWD). After a small craniotomy was made above the target region, 300 nl of the indicated AAV or AAV mixture was injected at each site using a microsyringe at 50 nl per min. The injection needle was maintained in place for 8–10 min after injection and then withdrawn slowly.

For basal forebrain (BF) injections, coordinates relative to bregma were AP, 0 mm; ML, ±1.35 mm; and DV, -5.1 mm from the dura. For dorsal striatum (DS) injections, coordinates were AP, +1.0 mm; ML, +2.4 mm; and DV, -2.65 mm from the dura. For hippocampal CA1 injections, coordinates were AP, -1.8 mm; ML, +1.2 mm; and DV, -1.3 mm from the dura.

For fiber photometry experiments, an optical fiber (200-μm core diameter, 0.39 NA; RWD) was implanted 0.1 mm above the injection site and secured to the skull using dental cement (3M). Mice recovered on a heating pad before being returned to their home cages and group-housed at two to three animals per cage. Recordings were performed 3 weeks after viral injection.

### *In vivo* fiber photometry recording and analysis

Fiber photometry recordings were performed using an FPS-410/470/561 system (Inper) controlled by Inper Signal software (v2.0.8). Green and red fluorescent sensors were excited using 470/5-nm and 561/5-nm LEDs, respectively, delivered at 10 Hz with a pulse duration of 15 ms and an optical power of 20–30 μW at the fiber tip. Fiber optic patch cords were pre-bleached with a high-power LED before each recording session to reduce background fluorescence.

Fluorescence signals were acquired using Inper Signal and analyzed offline using custom MATLAB scripts (R2022a; MathWorks), including subtraction of background autofluorescence. For sleep-wake recordings, baseline fluorescence (F_0_) was defined as the mean fluorescence during stable non-rapid eye movement (NREM) sleep epochs. For tail-suspension and treadmill-running experiments, *F*₀ was defined as the mean fluorescence during the pre-stimulus baseline period immediately preceding stimulus onset. ΔF/F_0_ was calculated as (F–F_0_)/F_0_. For comparisons across animals and recording sessions, ΔF/F_0_ values were converted to z-scores using the mean and standard deviation of the corresponding baseline period.

### EEG/EMG recording and sleep-wake state classification

To monitor sleep-wake states, EEG electrodes were implanted through craniotomies above the frontal and visual cortices, and EMG wires were inserted bilaterally into the neck muscles. The electrodes were connected to a miniature connector fixed to the skull with dental cement. During recordings, the connector was linked via a flexible cable to an electrical slip ring, allowing unrestricted movement while preventing cable twisting. EEG and EMG signals were amplified (NL104A, Digitimer), filtered (NL125/6, Digitimer), digitized using a Power1401 data acquisition interface (Cambridge Electronic Design), and recorded with Spike2 software (Cambridge Electronic Design) at a sampling rate of 1,000 Hz. Sleep-wake states were classified semi-automatically in 4-s epochs using AccuSleep and manually verified using a custom MATLAB GUI. Wakefulness was characterized by desynchronized cortical EEG activity and elevated EMG activity. NREM sleep was identified by synchronized EEG activity dominated by high-amplitude delta waves (0.5–4 Hz) and low EMG activity. REM sleep was characterized by prominent theta activity (6–9 Hz) together with minimal EMG activity.

### Tail suspension test

All behavioral tests were performed at least 1 h into the dark phase. Each mouse was gently lifted off the cage bottom six times for 60 seconds, with a minimum of 1 minute between each lift.

### Locomotion monitoring and running paradigm

Locomotor activity of head-fixed mice was monitored using a custom-built belt treadmill. Mice were head-fixed with their limbs resting on a spherical treadmill (15 cm diameter). The rotation of the treadmill was measured using a rotary encoder (US Digital #MA3-A10-250-N), which converted rotational velocity into an analog voltage signal (∼0–5 V). The signal was digitized at 1 kHz using a National Instruments USB-6001 multifunction data acquisition device with the MATLAB DAQ toolbox and custom-written software and converted to running speed at 10 Hz. Mice were habituated to treadmill running for 3 days prior to the experiment. Mice were first acclimated to the treadmill for 15 minutes. This was followed by a 1-h session of voluntary running. Subsequently, mice underwent three 3-min sessions of motor-driven forced running. Finally, to pharmacologically validate the rAdo1h sensor, an A_2A_R agonist JMF1907 (10 mg/kg) was administered intraperitoneally (i.p.).

### Fiber photometry combined with optogenetic manipulation

Adult CaMKIIα-Cre or VGLUT2-Cre mice were used for all experiments. To validate the sensors in combination with optogenetic manipulation, viruses were stereotaxically injected into the hippocampal CA1 region or the BF. Three weeks after viral injection, fiber photometry recordings were performed using an R801 tricolor multi-channel fiber photometry system (RWD Life Science). Fluorescence excitation was achieved using three LED/filter channels (405/10 nm, 470/40 nm, and 555/20 nm). Signals were acquired using a 10-Hz excitation protocol with 15 ms pulse duration at 20–30 μW power at the fiber tip. The 470 nm channel was used to monitor sensor activity, whereas the 405 nm isosbestic channel served to correct for motion artifacts and nonspecific signal fluctuations. For optogenetic experiments, a 635 nm red laser (5 mW at the fiber tip) was delivered through the same optical fiber to activate neurons expressing the red-shifted opsin ChrimsonR. Stimulation consisted of 20-Hz pulse trains lasting 4, 8, or 16 s, with a pulse width of 5 ms. Each stimulation was separated by a 300 s inter-stimulus interval, and trials were separated by at least 600 s. Photometry and optogenetic stimulation were synchronized to enable simultaneous recording and manipulation. For pharmacological experiments, VGLUT2-Cre mice received systemic administration of 5-ITU (1.6 mg/kg, i.p.), JMF1907 (10 mg/kg, i.p.), or SCH-58261 (10 mg/kg, i.p.) prior to recordings. After drug treatment, the same optogenetic stimulation protocols described above were applied to assess drug-dependent modulation of stimulation-evoked fluorescence signals. Prior to recording, mice were habituated to the setup for at least 30 min to reduce stress and ensure stable behavioral states.

### Two-photon *in vivo* imaging

Adult mice were anesthetized with isoflurane. The skin and skull covering the visual cortex were retracted, and 300 nl of the respective AAVs were injected into the visual cortex (AP: −3.08 mm relative to bregma; ML: +1.5 mm relative to bregma; DV: −0.5 mm from the dura). A 3 mm diameter round coverslip was used to replace the removed skull, and a metal recording chamber and EEG electrodes were securely attached. A stainless-steel head holder was attached to the skull to head-fix the animal and to reduce motion-induced artifacts during imaging. Two weeks post-virus injection, awake mice were habituated for ∼15 min in the treadmill-adapted imaging apparatus to minimize stress effects from head restraint. The visual cortex, at a depth of 100–200 μm below the pial surface, was imaged using Prairie View 5.5.64.100 software on a Bruker Ultima Investigator two-photon microscope equipped with the Spectra-Physics Insight X3 laser and a 16×/0.8 NA water immersion objective. For dual-color imaging, the fluorophores were excited at 920 nm and 1040 nm, with the corresponding emission collected through 525/70-nm and 595/50-nm emission filters, respectively. The sampling rate was set to approximately 1 Hz. An infrared video camera was positioned above the animal to capture behavioral correlates. Following ∼60 min of baseline recording after the saline injection, kainic acid (KA, 10 mg/kg, i.p.) was administered to elicit seizures. For image analysis, motion artifacts were first corrected using the EZcalcium^53^ motion correction algorithm. Subsequently, the fluorescence intensities of regions of interest (ROIs) were extracted using ImageJ software. The background-subtracted fluorescence intensity was then used to calculate the relative fluorescence change (ΔF/F_0_).

### Quantification and statistical analysis

Animals or cells were randomly assigned to experimental or control groups. Imaging data were processed using ImageJ (NIH) and custom-written MATLAB scripts (R2020b). Data visualization and statistical analysis were performed using OriginPro (OriginLab). Unless otherwise indicated, summary data are presented as mean ± s.e.m. and each data point represents one sample. Signal-to-noise ratio (SNR) was calculated as the peak fluorescence response divided by the standard deviation of baseline fluorescence fluctuations. Comparisons between two groups were performed using two-sided Student’s *t*-test, Mann-Whitney *U* test, paired *t*-test, or Wilcoxon signed-rank test, as appropriate. Multiple-group comparisons were performed using two-way ANOVA followed by Bonferroni’s multiple-comparisons test. Correlations were assessed using Pearson correlation analysis. Statistical significance was defined as ns, not significant (*P* ≥ 0.05); \**P* < 0.05; \*\**P* < 0.01; and \*\*\**P* < 0.001.

## Acknowledgements

We thank Drs. Lei Ma, Hui Dong, Jingchao Zhang, Wu and Li Lab members for critical reading of the manuscript. We are grateful to Dr. Miao Jing at the Chinese Institute for Brain Research and Dr. Shu Xie at Peking University for technical support with two-photon imaging; to Dr. Lei Ma and Liang Chen at Nanjing Medical University for help with the treadmill setup; and to Ms. Liqin Fu at the National Center for Protein Sciences, Peking University, and Dr. Lei Su at the Institute of Microbiology, Chinese Academy of Sciences, for technical support with confocal imaging. This work was financially supported by the National Natural Science Foundation of China (32422031 to Z.W. and 323B2032 to B.L.), the National Key R&D Program of China (2022YFE0108700 to Y.L.), the STI2030-Major Projects (2022ZD0208300 to Z.W.), the CAS Project for Young Scientists in Basic Research (YSBR-126 to Z.W.), and the Xiaomi Foundation (to Z.W.).

## Author contributions

Z.W. and YL.L. conceived the project and designed the experiments. B.L., W.T., and K.H. developed and optimized the sensors. B.L. and P.H. characterized the sensors in cell cultures under the supervision of Z.W. and YL.L. YM.L. and Z.B. performed the surgery and *in vivo* two-photon imaging experiments in head-fixed mice under the supervision of Z.W. YM.L., J.W. (Capital Medical University), Z.M., K.C., Y.H., and F.L. performed the fiber photometry recordings in freely moving mice under the supervision of Z.W. Z.W., YM.L., and B.L. wrote the manuscript. All authors contributed to data analysis and interpretation.

---

**Extended Data Fig. 1.**
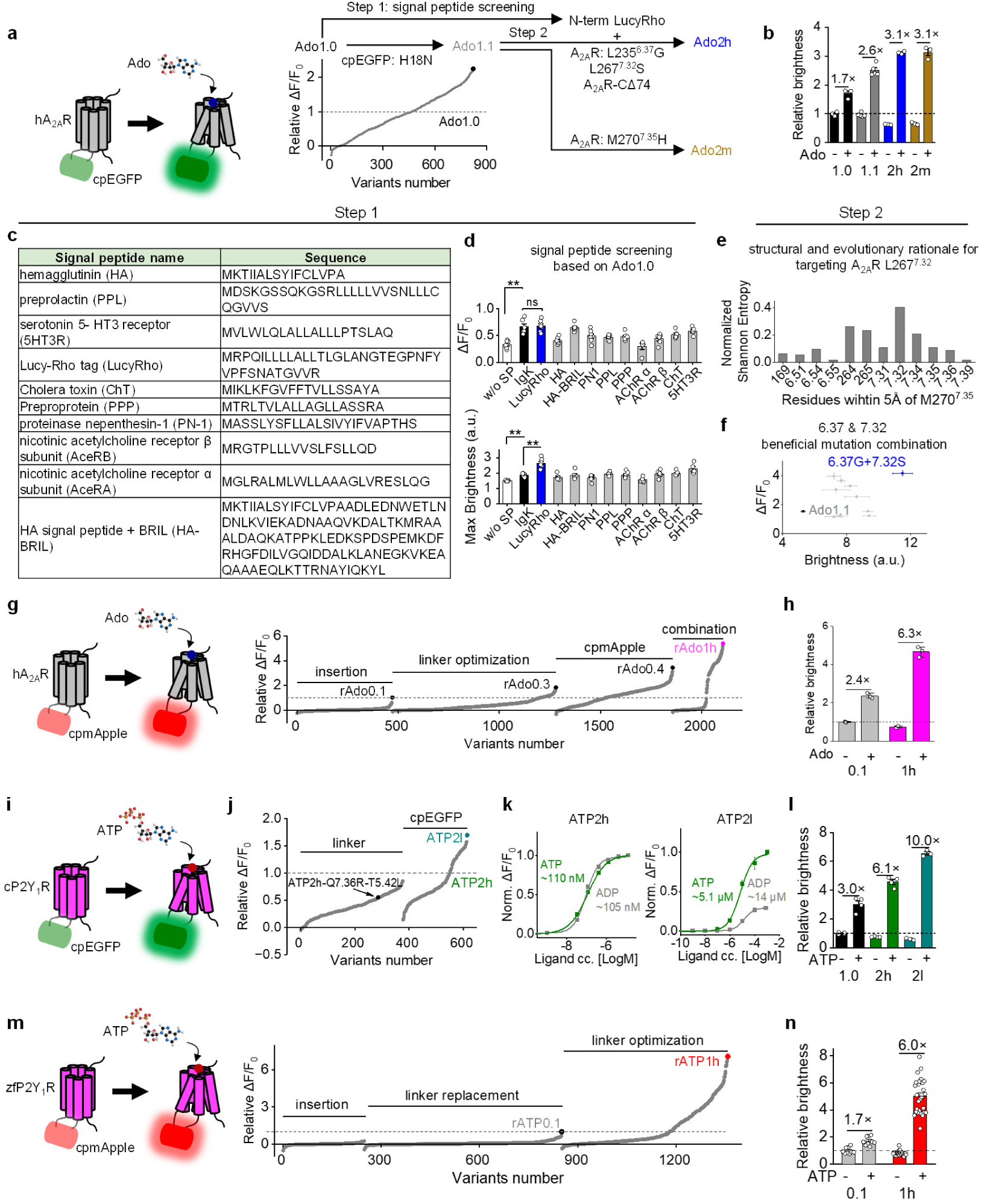
Engineering trajectories of next-generation adenosine and ATP sensors. **a**, Schematic of the green Ado sensor architecture and screening trajectory from Ado1.0 to Ado1.1, Ado2m, and Ado2h. Ado1.1 contains the cpEGFP mutation H18N; Ado2m was generated by introducing M270^7.35^H into the A_2A_R scaffold; and Ado2h incorporates the Lucy-Rho N-terminal tag, a 74-amino-acid C-terminal truncation, L235^6.37^G, and L267^7.32^S. The plot shows the relative ΔF/F₀ of the screened variants. **b**, Relative fluorescence brightness of Ado1.0, Ado1.1, Ado2h, and Ado2m before (−) and after (+) Ado application. **c**, Names and amino-acid sequences of the N-terminal signal peptides tested during green Ado sensor optimization. Abbreviations shown in parentheses are used in **d**. **d**, Screening of the indicated signal peptides using Ado1.0 as the sensor scaffold. IgK denotes Ado1.0 containing its original immunoglobulin κ signal peptide; w/o SP denotes Ado1.0 with the N-terminal signal peptide removed. Top, peak Δ*F*/*F*₀, *P* = 0.0012 (w/o SP vs. IgK), *P* = 0.84 (IgK vs. LucyRho); bottom, maximum fluorescence brightness, *P* = 0.0012 (w/o SP vs. IgK), *P* = 0.0012 (IgK vs. LucyRho). **e**, Sequence-conservation analysis of A_2A_R residues located within 5 Å of M270^7.35^. **f**, Comparison of the affinity index and ΔF/F₀ for Ado1.1 variants carrying individual or combined mutations at A_2A_R positions L235^6.37^ and L267^7.32^. The affinity index was calculated as the ratio of ΔF/F₀ evoked by 30 nM NECA to that evoked by 100 μM Ado. **g**, Schematic of the red Ado sensor architecture and screening trajectory through insertion-site screening, linker optimization, cpmApple optimization, and combination of selected mutations. The indicated intermediates are rAdo0.1, rAdo0.3, and rAdo0.4, followed by rAdo1h. **h**, Relative fluorescence brightness of rAdo0.1 and rAdo1h before (−) and after (+) Ado application. **i**, Schematic of the green ATP sensor comprising cpEGFP inserted into the chicken P2Y_1_R. **j**, Screening trajectory from ATP2h-Q^7.36^R-T^5.42^L to ATP2l through linker and cpEGFP optimization. The plot shows the relative ΔF/F_0_ of the screened variants. **k**, ATP and ADP concentration-response curves for ATP2h and ATP2l in cultured neurons. **l**, Relative fluorescence brightness of ATP1.0, ATP2h, and ATP2l before (−) and after (+) ATP application. **m**, Schematic of the red ATP sensor comprising cpmApple inserted into the zebra finch P2Y_1_R, together with the screening trajectory through insertion-site screening, linker replacement, and linker optimization. The indicated intermediate and final sensors are rATP0.1 and rATP1h, respectively. **n**, Relative fluorescence brightness of rATP0.1 and rATP1h before (−) and after (+) ATP application. For **b**,**h**,**l** and **n**, fluorescence was measured in HEK293T cells before and after application of 100 µM Ado (**b** and **h**) or 100 µM ATP (**l** and **n**); *n* = 3–4 wells per sensor in 96-well plates. Relative brightness was normalized to the ligand-free baseline brightness of Ado1.0, rAdo0.1, ATP1.0, and rATP0.1 in **b**,**h**,**l** and **n**, respectively. Data are presented as mean ± s.e.m. Each data point represents one sample. Mann-Whitney *U* test; ns, not significant, \*\**P <* 0.01.

**Extended Data Fig. 2.**
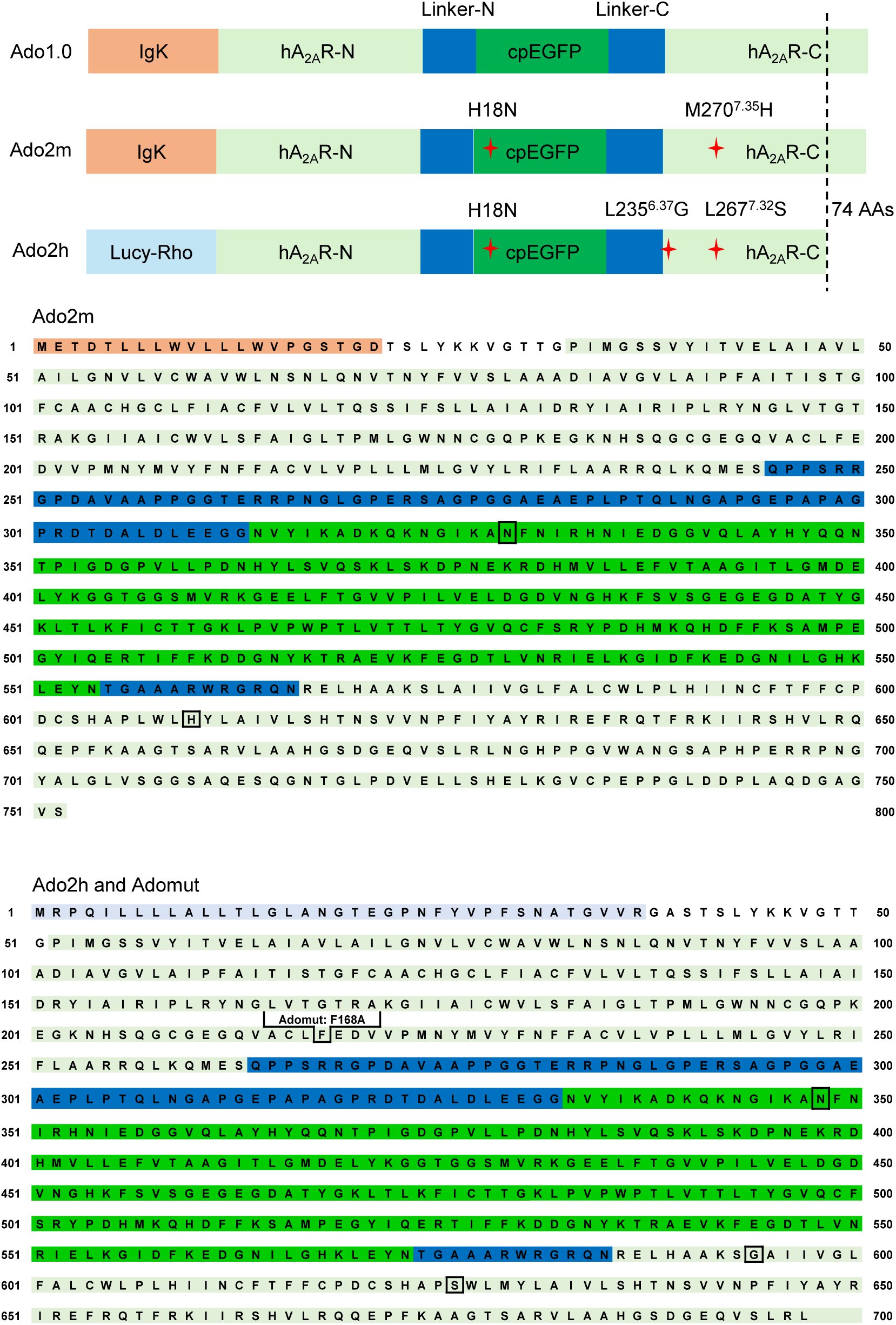
Full amino acid sequence of the Ado2m, Ado2h and Adomut sensors. The IgK leader sequence, Lucy-Rho leader sequence, cpEGFP moiety, N-terminal and C-terminal portions of the human A_2A_R (hA_2A_R) are indicated. A C-terminal truncation of 74 amino acids has been introduced into Ado2h. The asterisk represents the location of a point mutation. Orange indicates the IgK leader sequence, pale blue indicates the Lucy-Rho leader sequence, light green indicates the hA_2A_R, dark blue indicates the N- and C-terminal linkers, and green indicates cpEGFP. Red asterisks in the schematics and boxed residues in the sequences indicate mutations incorporated into the indicated sensor or ligand-insensitive mutant sensor.

**Extended Data Fig. 3.**
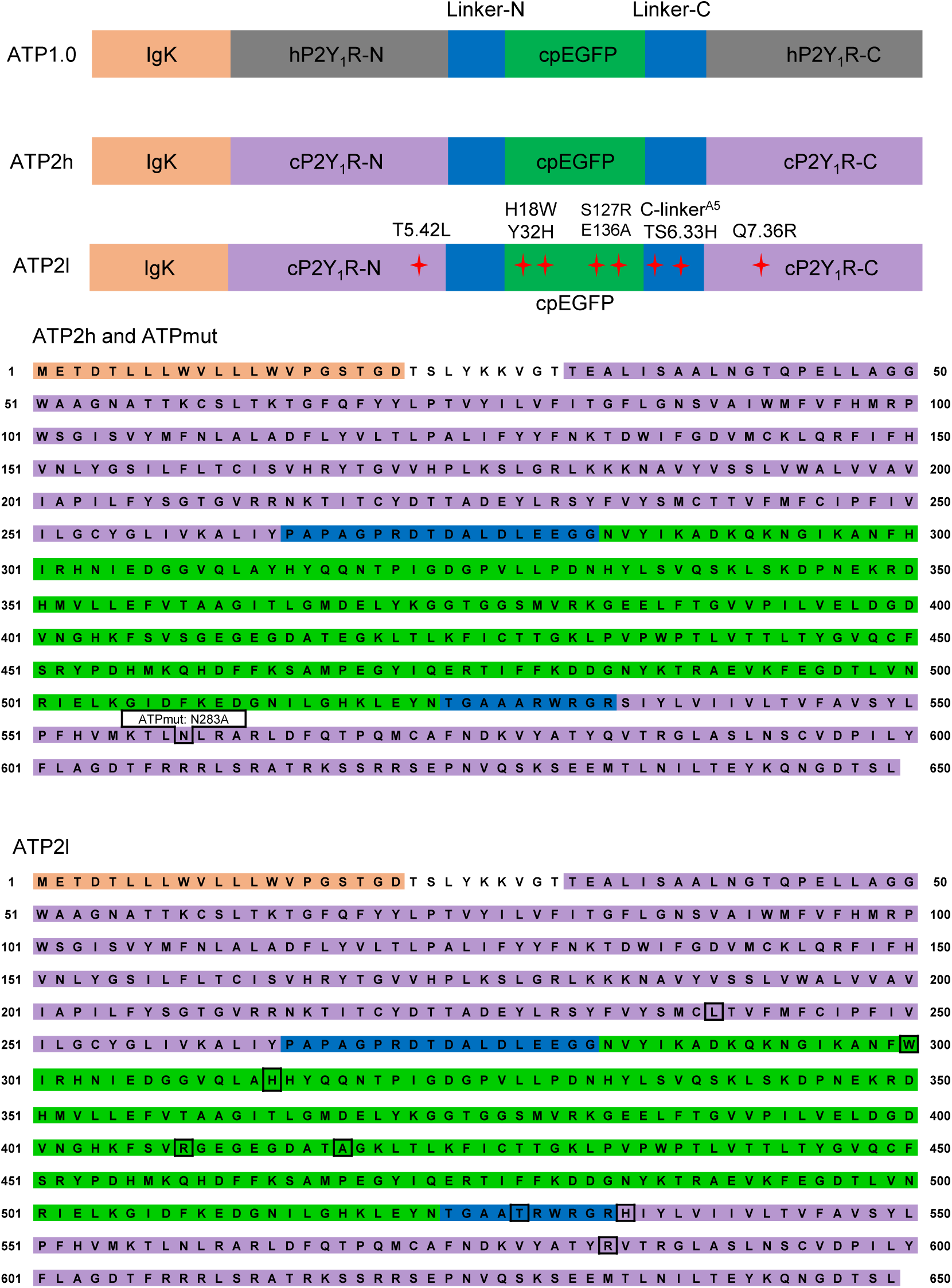
Full amino acid sequence of the ATP2h, ATP2l and ATPmut sensors. The IgK leader sequence, cpEGFP moiety, N-terminal and C-terminal portions of the human or chicken P2Y_1_R (hP2Y_1_R or cP2Y_1_R) are indicated. The asterisk represents the location of a point mutation. Orange indicates the IgK leader sequence, grey indicates the hP2Y_1_R, purple indicates the cP2Y_1_R, dark blue indicates the N- and C-terminal linkers, and green indicates cpEGFP. Red asterisks in the schematics and boxed residues in the sequences indicate mutations incorporated into the indicated sensor or ligand-insensitive mutant.

**Extended Data Fig. 4.**
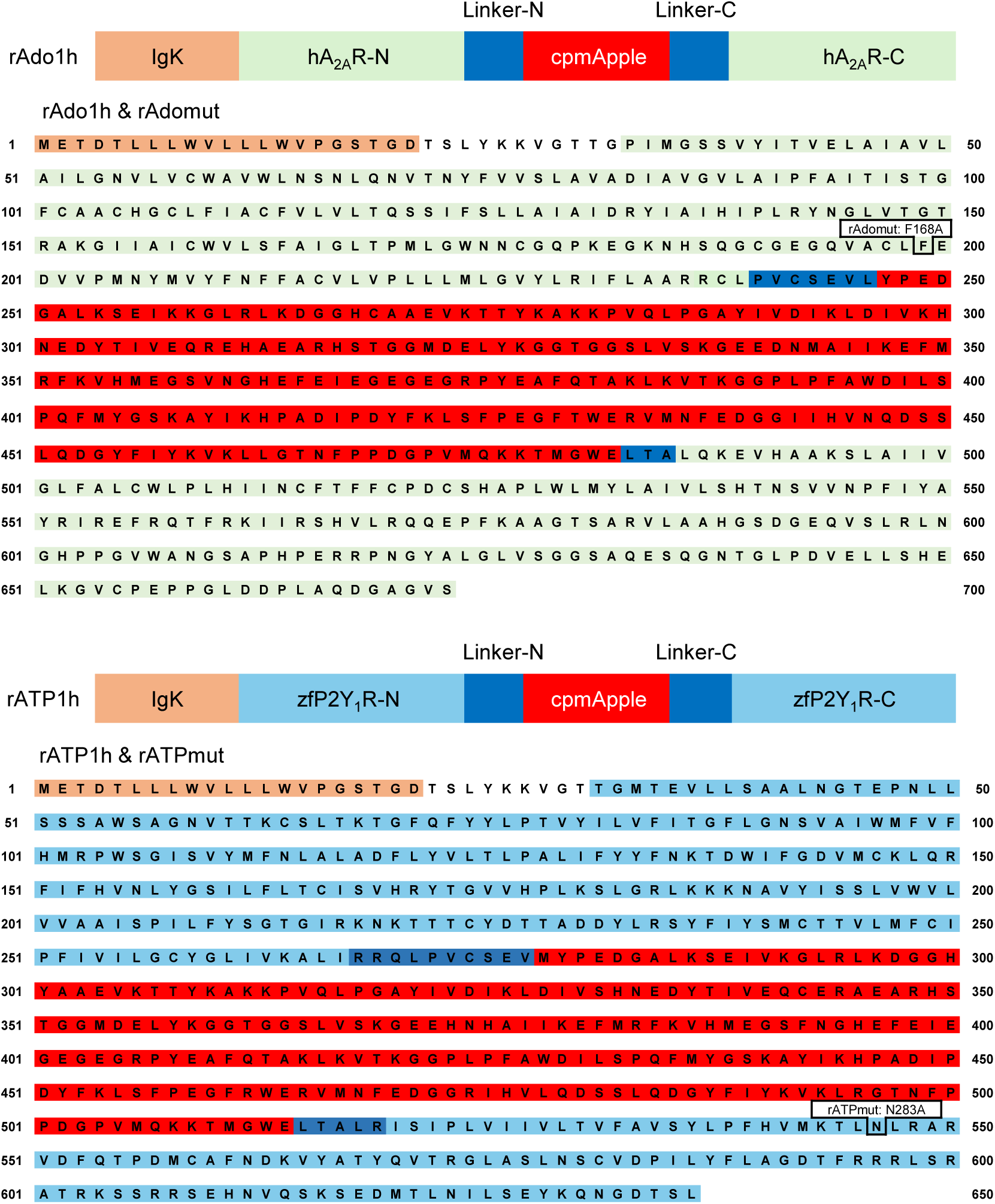
Full amino acid sequence of the rAdo1h, rAdomut, rATP1h and rATPmut sensors. The IgK leader sequence, cpmApple moiety, N-terminal and C-terminal portions of the hA_2A_R and zebra finch P2Y_1_R (zfP2Y_1_R) are indicated. Orange indicates the IgK leader sequence, light green indicates the hA_2A_R, light blue indicates the zfP2Y_1_R, dark blue indicates the linkers, and red indicates cpmApple. Boxed residues indicate mutations incorporated into the indicated sensor or ligand-insensitive mutant.

**Extended Data Fig. 5.**
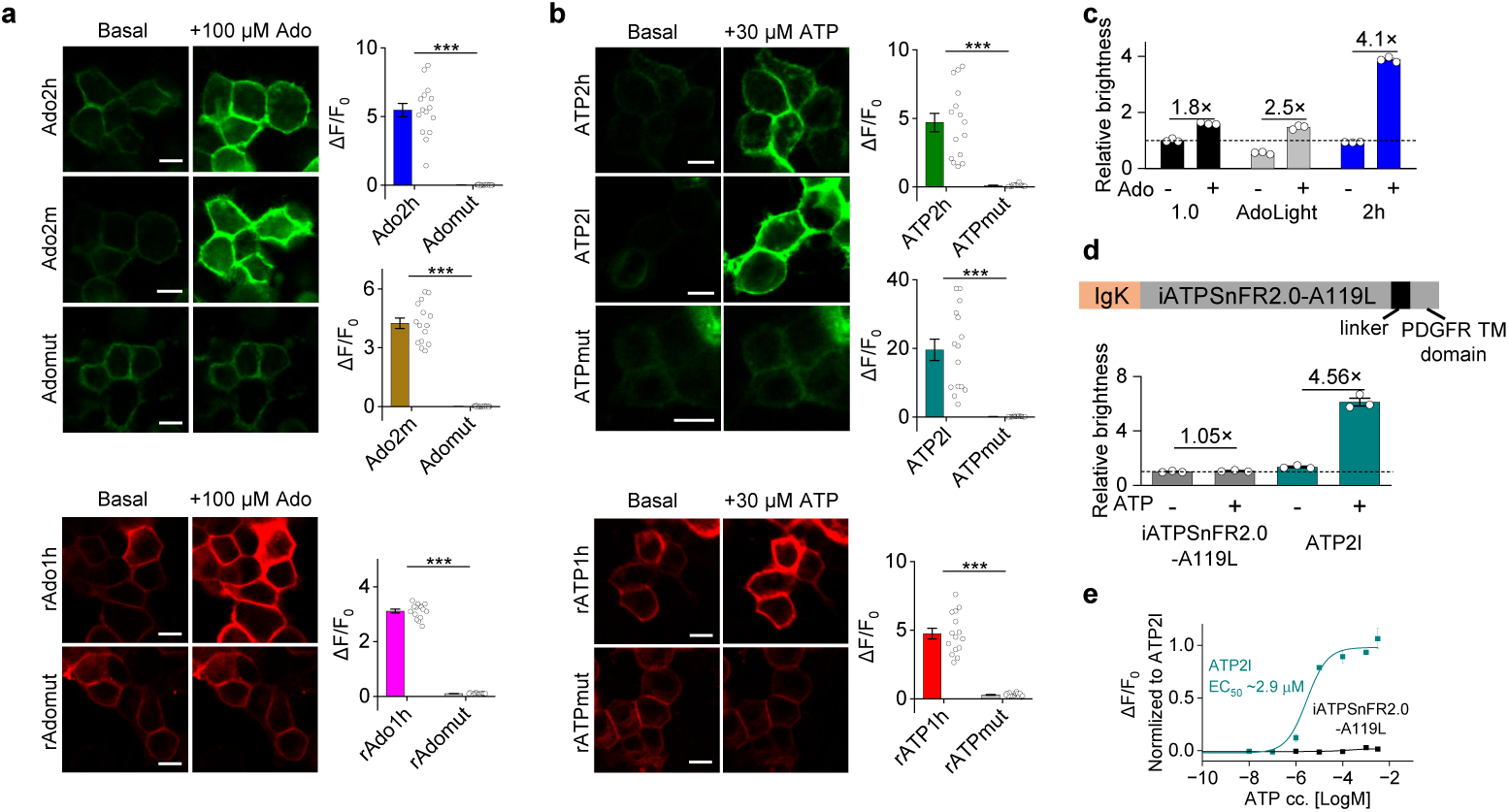
Performance of mutants and other adenosine or ATP sensors. **a**, Representative images (left) and group summary of maximal ΔF/F_0_ in response to 100 μM Ado (right) of indicated sensors in HEK293T cells. Scale bar, 10 μm. *P <* 0.001 (Ado2h vs. Adomut, Ado2m vs. Adomut and rAdo1h vs. rAdomut). *n* = 15 cells from 3 coverslips per group. The same Adomut control data are displayed in the Ado2h-versus-Adomut and Ado2m-versus-Adomut comparisons. **b**, Representative images (left) and group summary of maximal ΔF/F_0_ in response to 30 μM ATP (right) of indicated sensors in HEK293T cells. Scale bar, 10 μm. *P <* 0.001 (ATP2h vs. ATPmut, ATP2l vs. ATPmut and rATP1h vs. rATPmut). *n* = 15/3 for each group. The same ATPmut control data are displayed in the ATP2h-versus-ATPmut and ATP2l-versus-ATPmut comparisons. **c,** Relative fluorescence brightness of Ado1.0, AdoLight, and Ado2h in HEK293T cells before (−) and after (+) application of 100 µM Ado. Values were normalized to the ligand-free brightness of Ado1.0. Numbers indicate the +Ado/−Ado fold change. **d,** Top, schematic of iATPSnFR2.0-A119L containing the IgK signal peptide, iATPSnFR2.0-A119L module, linker, and PDGFR transmembrane (TM) domain. Bottom, relative fluorescence brightness of iATPSnFR2.0-A119L and ATP2l before (−) and after (+) application of 1 mM ATP. Values were normalized to the ligand-free brightness of iATPSnFR2.0-A119L. Numbers indicate the +ATP/−ATP fold change. **e,** ATP concentration-response curves for ATP2l and iATPSnFR2.0-A119L in HEK293T cells. Responses were normalized to the maximal ATP2l response. Data are presented as mean ± s.e.m. Each data point represents one sample. Student’s *t*-test; *** *P <* 0.001.

**Extended Data Fig. 6.**
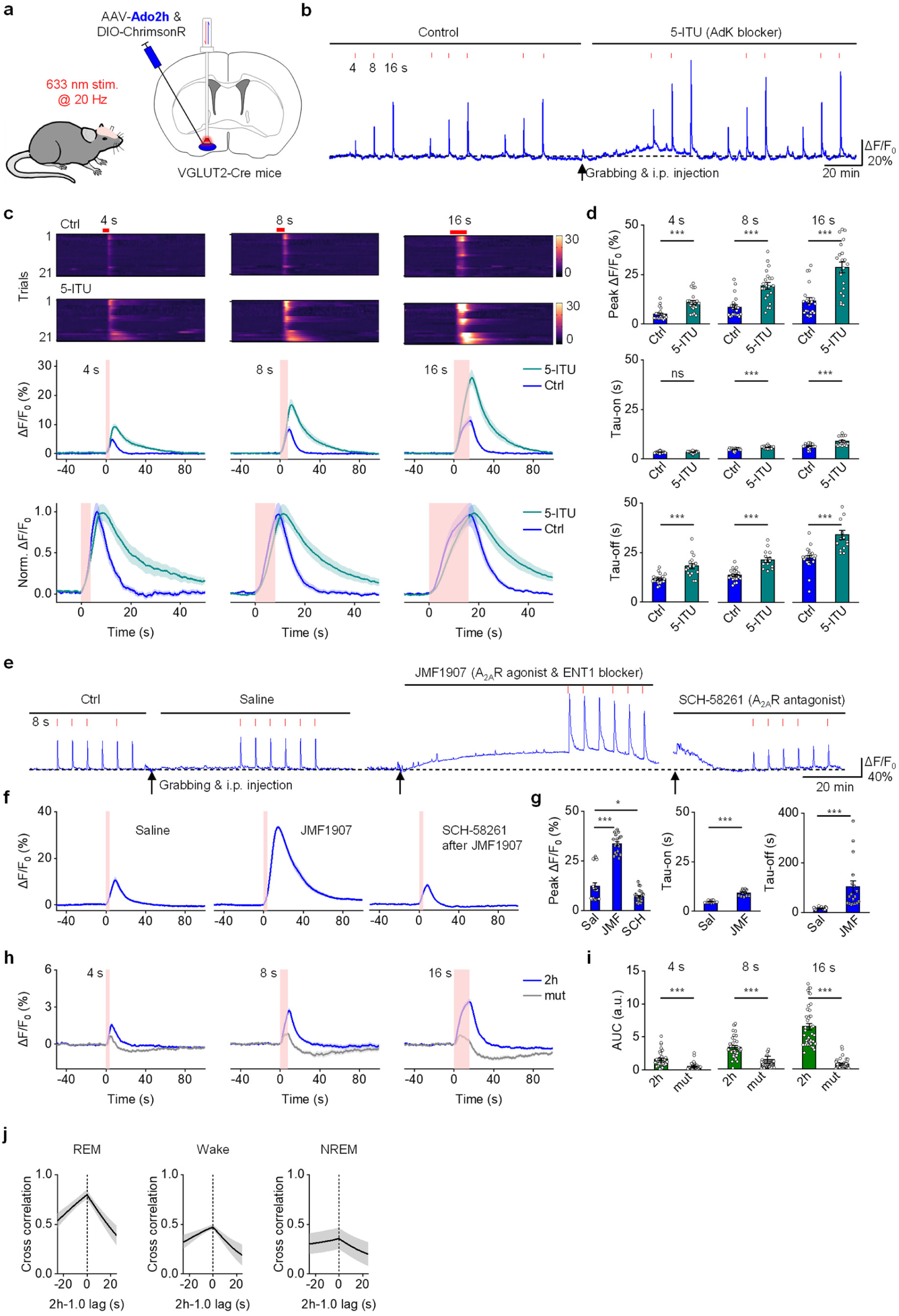
Detection of Ado release evoked by optogenetics in freely moving mice. **a,** The experimental design described the strategy for expressing Ado2h and recorded fluorescence changes during optical stimulation in the BF of VGLUT2-Cre mice. **b,** Representative traces show optogenetically stimulated fluorescence changes in Ado2h in the BF before (baseline left) and after an i.p. injection of the AdK blocker 5-ITU (1.6 mg/kg, right). The vertical red tick marks indicate optogenetic stimulation lasting 4, 8, or 16 seconds at 20 Hz. **c,** Ado2h signals aligned to optogenetic stimulation lasting 4, 8, or 16 seconds before (blue) and after an i.p. injection of 5-ITU (green) at 20 Hz for all mice. (Top) Heat map plot of 21 individual trials from 3 mice; (middle) averaged signal; (bottom) normalized averaged signal. The red shaded area and red line indicate the application of opto-stimulation. **d,** Group summary of peak response of Ado2h (top), rise time (Tau-on), and decay time (Tau-off) before (blue) and after an i.p. injection of 5-ITU (green) during the indicated optogenetic stimulation durations (4, 8, or 16 s). *n* = 20–21 trials in 3 mice per group. For peak response, *P <* 0.001 for all durations. For Tau-on, *P =* 0.88, *P* < 0.001, and *P* < 0.001 for stimulation durations of 4, 8, and 16 s, respectively; for Tau-off, *P <* 0.001 for all durations. **e,** Representative traces show the optogenetically stimulated fluorescence changes in Ado2h in the BF at baseline and after injection of saline. Further traces illustrate changes following an i.p. injection of the A_2A_R agonist and ENT1 blocker JMF1907 (10 mg/kg), and after an i.p. injection of the A_2A_R antagonist SCH-58261 (10 mg/kg). Red tick marks indicate 8-s optical stimulation at 20 Hz. **f,** Averaged Ado2h responses following saline, JMF1907, and SCH-58261 injections. The red bar indicates an 8-s stimulation at 20 Hz. **g,** Group summaries of peak ΔF/F₀ after saline, JMF1907, or SCH-58261 injection (left), and Tau-on (middle) and Tau-off (right) after saline or JMF1907 injection. For peak ΔF/F₀, *P* < 0.001 for saline versus JMF1907 and *P* = 0.03 for saline versus SCH-58261. **h,** Average traces showing changes in the sensors Ado2h (blue) and Adomut (grey) in response to 4-, 8-, or 16-second optical stimulation. The red-shaded area indicates the period of optical stimulation. **i,** AUC of Ado2h and Adomut responses during the indicated stimulation durations. *P* < 0.001 for Ado2h versus Adomut at each duration; *n* = 34–40 trials from 3 mice per group. **j,** Cross-correlation of Ado1.0 and Ado2h fluorescence signals during each vigilance state. Data are presented as mean ± s.e.m. Each data point represents one sample. Paired two-tailed Student’s *t*-tests were used for within-mouse drug comparisons in **d** and **g**; unpaired two-tailed Student’s *t*-tests were used for comparisons between Ado2h and Adomut in **i**; ns, not significant, \**P* < 0.05, \*\*\**P* < 0.001.

**Extended Data Fig. 7.**
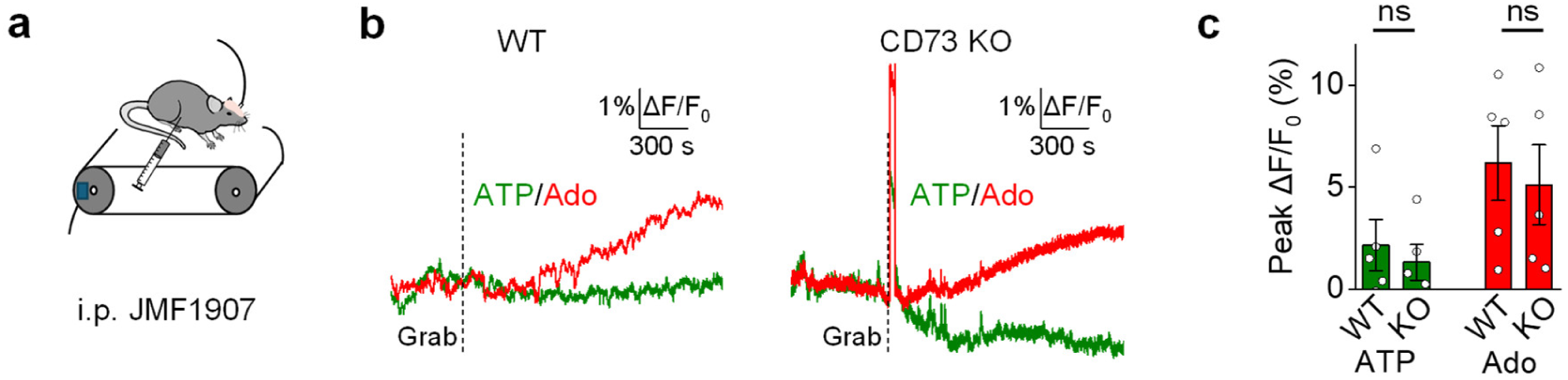
Detection of JMF1907-induced rAdo1h responses in WT and CD73 KO mice. **a,** Schematic of the experimental setup for fiber photometry recording of ATP2h and rAdo1h sensors after injection of JMF1907. **b,** Average traces of the change in fluorescence from a WT (left) and CD73 KO (right) mouse. The dashed line indicates the injection of JMF1907. **c,** Group summary of the peak change in fluorescence of indicated sensors after JMF1907 injection. *n* = 5 trials from 5 mice per group. *P* = 0.69 for ATP2h and *P* > 0.99 for rAdo1h. Data are presented as mean ± s.e.m. Each data point represents one sample. Mann-Whitney *U* test; ns, not significant.

**Extended Data Fig. 8.**
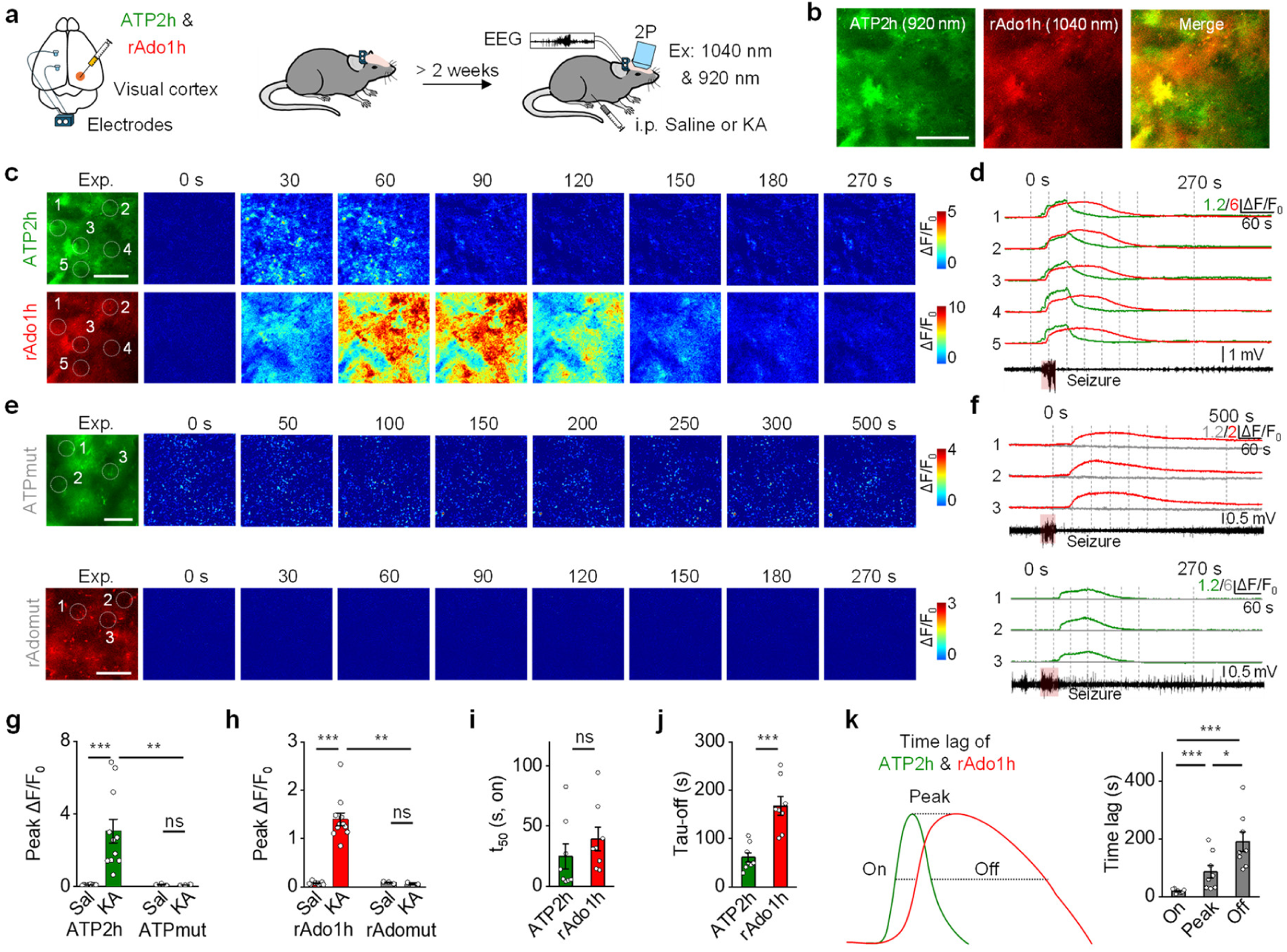
*In vivo* dual-color two-photon imaging of ATP2h and rAdo1h in a mouse seizure model. **a,** Schematic illustration depicting virus injection of ATP2h and rAdo1h and head-fixed two-photon imaging in the mouse visual cortex (V1) during kainic acid (KA)-induced seizure. **b,** Representative *in vivo* two-photon image of V1 showing ATP2h and rAdo1h fluorescence. Scale bar, 100 μm. **c,** Representative images show ATP2h (top) and rAdo1h (bottom) expression and responses during seizure. Scale bar, 50 μm. **d**, Traces measured at indicated ROIs (white circles) during seizure. **e,f,** Similar to **c,d** except that the mice were injected with the ATPmut virus (top) or rAdomut virus (bottom). **g,** Group summary of the peak fluorescence changes of ATP2h and ATPmut under the indicated treatments; *P <* 0.001 (saline vs. KA of ATP2h), *P =* 0.70 (saline vs. KA of ATPmut) and *P =* 0.006 (ATP2h vs. ATPmut after KA injection). *n* = 11 and 3 mice for ATP2h and ATPmut, respectively. **h,** Group summary of the peak fluorescence changes of rAdo1h and rAdomut under the indicated treatments; *P <* 0.001 (saline vs. KA of rAdo1h), *P =* 0.20 (saline vs. KA of rAdomut) and *P =* 0.006 (rAdo1h vs. rAdomut after KA injection). *n* = 11 and 3 mice for rAdo1h and rAdomut, respectively. **i,** Group summary of the rise t_50_ of ATP2h and rAdo1h signals during seizure. *P =* 0.13; *n* = 8 mice per group. **j,** Group summary of the decay time constant of ATP2h and rAdo1h signals during epileptic seizure; *P <* 0.001. *n* = 8 mice per group. **k,** Schematic diagram (left) and group summary (right) of time lag of ATP2h and rAdo1h signals during seizure; *P <* 0.001 (on vs. peak), *P <* 0.001 (on vs. off) and *P =* 0.04 (peak vs. off). *n* = 8 mice per group. Data are presented as mean ± s.e.m. Each data point represents one sample. Mann-Whitney *U* test; ns, not significant, \**P* < 0.05, \*\**P* < 0.01, \*\*\**P* < 0.001.

**Extended Data Fig. 9.**
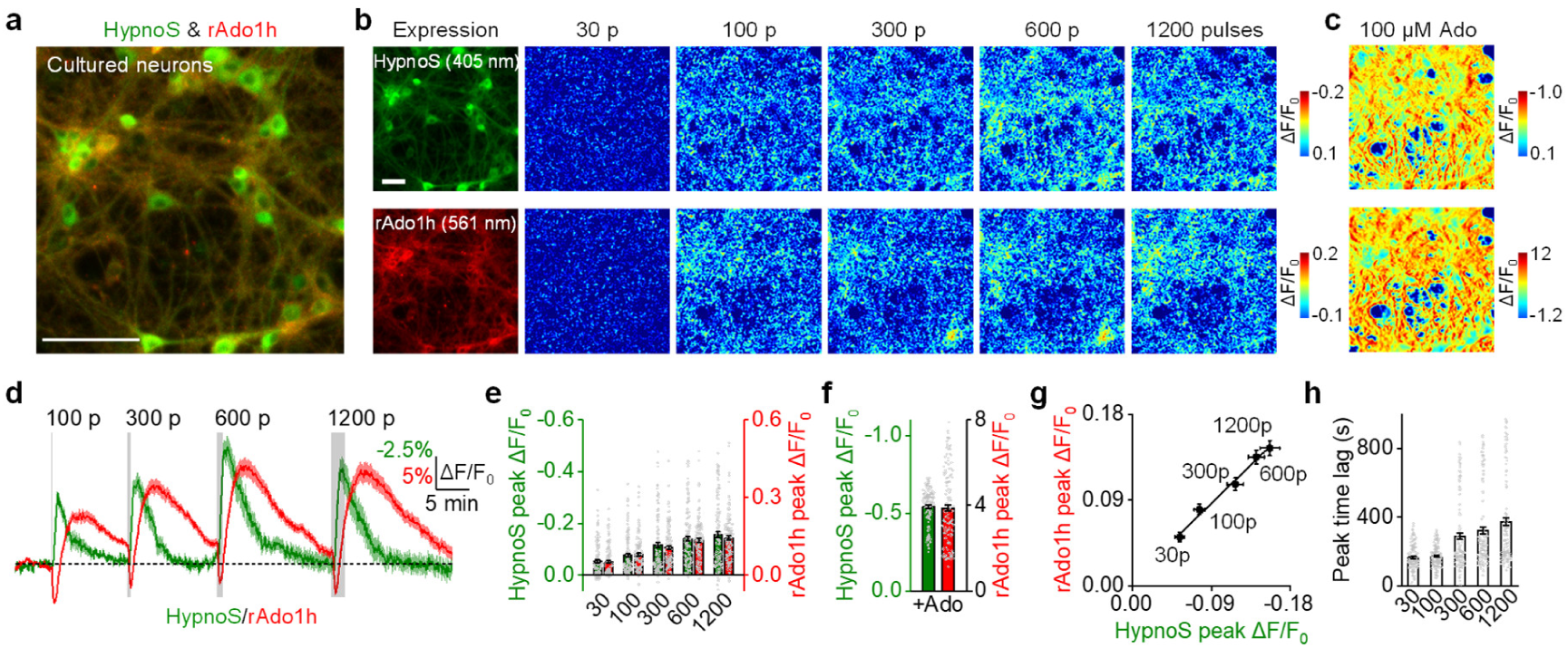
*In vitro* recording of HypnoS and rAdo1h responses in cultured neurons. **a,** Representative images showing the expression of HypnoS and rAdo1h in neurons. Scale bar, 100 μm. **b,** Representative images show HypnoS (top) and rAdo1h (bottom) expression and responses to electrical stimulation. Scale bar, 50 μm. **c,** Representative images of HypnoS (top) and rAdo1h (bottom) response to application of 100 μM Ado. Scale bar, 50 μm. **d,** Average traces of the change in HypnoS and rAdo1h fluorescence. The gray-shaded area indicates the application of electrical stimulation. **e,** Group summary of the peak fluorescence changes of HypnoS and rAdo1h to indicated electrical stimulus. *n* = 139–148 neurons per group. **f,** Group summary of the peak fluorescence changes of HypnoS and rAdo1h to application of 100 μM Ado. *n* = 148 neurons per group. **g,** Correlation between the magnitudes of peak HypnoS and rAdo1h fluorescence responses across five electrical-stimulation intensities. Pearson’s *r*(3) = 0.994, two-sided *P* = 0.000571, 95% CI [0.907, 1.000]. Each point represents the mean response at one stimulation intensity; *n* = 5 stimulation intensities, derived from 139–148 neurons per condition from 3 independent cultures. **h,** Group summary of peak time lag of HypnoS and rAdo1h signals during electrical stimulation. *n* = 139–148 neurons per group. Data are presented as mean ± s.e.m. Each data point represents one sample.

