## Supplementary Information for "An expanded GRAB sensor palette for multiplexed imaging of purinergic dynamics *in vivo*"

**Supplementary Notes**

**Green adenosine sensor engineering**

To improve the first-generation green adenosine sensor Ado1.0^1^, we first screened 14 positions in cpEGFP and identified H18N as the only beneficial substitution. The resulting intermediate was designated Ado1.1. We next pursued separate medium- and high-affinity branches (**Extended Data Fig. 1a**). Introducing M270^7.35^H into A_2A_R yielded the medium-affinity sensor Ado2m.

For the high-affinity branch, we used a two-stage optimization strategy comprising N-terminal signal-sequence optimization and GPCR scaffold optimization. First, we optimized the N terminus using Ado1.0 as the screening backbone. We tested a panel of signal sequences **(Extended Data Fig. 1c)** and quantified both peak ΔF/F_0_ and maximum fluorescence brightness for each candidate. This screen identified the hybrid Lucy-Rho motif, which combines a cleavable leucine-rich signal peptide with a non-cleavable rhodopsin-derived tag^2^, as the preferred replacement for the original IgK signal sequence **(Extended Data Fig. 1d)**.

Second, we optimized the A_2A_R scaffold. Screening positions implicated in GPCR activation^3^ identified L235^6.37^G as beneficial. Although M270^7.35^H increased sensor response, it also reduced apparent affinity^4^. We therefore sought a nearby position whose mutation might enhance response through local conformational coupling while better preserving apparent affinity. We aligned A_2A_R orthologs curated in GPCRdb^5^ and examined residues surrounding M270^7.35^. L267^7.32^, which lies within 5 Å of M270^7.35^, was less conserved across species than neighboring residues and was therefore selected for mutagenesis (**Extended Data Fig. 1e**). Using Ado1.1-L235^6.37^G as the screening backbone, we performed site-directed mutagenesis at L267^7.32^ and evaluated the resulting variants using ΔF/F₀ and an affinity index, defined as the ratio of the ΔF/F₀ evoked by 30 nM NECA to that evoked by 100 μM Ado. This screen identified L267^7.32^S as the preferred substitution. We additionally truncated the final 74 amino acids of the A_2A_R C terminus, following a strategy adapted from A_2A_R crystallography constructs^6,7^.

The resulting high-affinity sensor, Ado2h, combined the Lucy-Rho N-terminal tag with the engineered A_2A_R scaffold containing L235^6.37^G, L267^7.32^S, and CΔ74 **(Extended Data Fig. 1a,f**). After Ado application, relative fluorescence brightness increased 1.7-, 2.6-, 3.1-, and 3.1-fold for Ado1.0, Ado1.1, Ado2h, and Ado2m, respectively (**Extended Data Fig. 1b**).

**Red adenosine sensor engineering**

To construct a red adenosine sensor, we inserted cpmApple from GRAB_rDA1m_^8^, spanning PVCSE to LTV, into the third intracellular loop of wild-type human A_2A_R. Combinatorial screening paired N-terminal insertion sites from A203^5.64^ to S213^5.74^ with C-terminal insertion sites from P217 to V229^6.31^. We selected the variant with cpmApple inserted between L208^5.69^ and L225^6.27^ and designated it rAdo0.1. Optimization of the insertion-adjacent regions retained Q207^5.68^C in A_2A_R and M7L and V246A in cpmApple. We then randomized 17 cpmApple positions based on the evolutionary trajectories of mApple-related red fluorescent proteins^8,9^. Screening and combining beneficial substitutions retained Y30C, S60K, C71R, R73H, R191T, and R224L. Sequential insertion-site, linker, and cpmApple optimization produced rAdo0.1, rAdo0.3, and rAdo0.4 as intermediates before the final sensor rAdo1h (**Extended Data Fig. 1g**). After Ado application, relative fluorescence brightness increased 2.1-fold for rAdo0.1 and 6.1-fold for rAdo1h (**Extended Data Fig. 1h**).

**Green ATP sensor engineering**

To construct next-generation green ATP sensors, the cpEGFP module from ATP1.0^10^ was inserted into the chicken P2Y_1_R (UniProtKB P34996)^11^. Insertion-site screening identified a sensor with cpEGFP inserted between Y235^5.67^ and S247^6.33^, which was named ATP2h (**Extended Data Fig. 1i**). To reduce its apparent affinity, we screened residues surrounding the ligand-binding pocket^12,13^ and other positions reported to affect P2Y_1_R EC_50_^14^. This screen serendipitously identified Q^7.36^R, which improved ATP-over-ADP selectivity; adding T^5.42^L further reduced apparent ATP affinity. Starting from ATP2h-Q^7.36^R-T^5.42^L, sequential linker and cpEGFP optimization generated ATP2l (**Extended Data Fig. 1j**). In addition to Q^7.36^R and T^5.42^L, ATP2l retained H18W, Y32H, S127R, and E136A in cpEGFP, A5T in the C-terminal linker, and S247^6.33^H in the receptor. In cultured neurons, concentration-response measurements showed that ADP had a threefold lower apparent affinity than ATP and 0.28 times its maximal efficacy, supporting ATP-over-ADP selectivity (**Extended Data Fig. 1k**). After ATP application, relative fluorescence brightness changed 3.0-, 6.1-, and 10.0-fold for ATP1.0, ATP2h, and ATP2l, respectively (**Extended Data Fig. 1l**).

**Red ATP sensor engineering**

To develop a red ATP sensor, we first inserted cpmApple from GRAB_rDA1m_ into the third intracellular loop of zebra finch P2Y_1_R (UniProtKB H0ZM60)^11^. All variants in this screen decreased fluorescence after ligand application, suggesting inefficient coupling between receptor activation and cpmApple. We therefore repeated insertion-site screening using an optimized cpmApple module from GRAB_r5HT1.0_^15^, flanked by the RRQL and LQ linkers derived from rAdo0.1. A variant with RRQL-cpmApple-LQ inserted between I235^5.66^ and K247^6.32^ increased fluorescence and was designated rATP0.1. Further optimization retained V246A in cpmApple, a Q-to-R substitution at the second position of the C-terminal linker, K247^6.32^I, and Y250^6.35^P, yielding rATP1h (**Extended Data Fig. 1m**). After ATP application, relative fluorescence brightness changed 1.7-fold for rATP0.1 and 6.0-fold for rATP1h (**Extended Data Fig. 1n**). Full sequences of the optimized green adenosine, green ATP, and red sensors are provided in **Extended Data Figs. 2–4**, respectively; ligand-insensitive controls are characterized in **Extended Data Fig. 5**.
